# Estimating multiplicity of infection, allele frequencies, and prevalences accounting for incomplete data

**DOI:** 10.1101/2023.06.01.543300

**Authors:** Meraj Hashemi, Kristan A. Schneider

**Affiliations:** Department of Applied Computer- and Biosciences, University of Applied Sciences Mittweida, Mittweida, Germany

**Keywords:** social entrainment, daily activity patterns, circadian rhythms, spotted hyenas, behavioural classification, biologging

## Abstract

**Background:** Molecular surveillance of infectious diseases allows the monitoring of pathogens beyond the granularity of traditional epidemiological approaches and is well-established for some of the most relevant infectious diseases such as malaria. The presence of genetically distinct pathogenic variants within an infection, referred to as multiplicity of infection (MOI) or complexity of infection (COI) is common in malaria and similar infectious diseases. It is an important metric that scales with transmission intensities, potentially affects the clinical pathogenesis, and a confounding factor when monitoring the frequency and prevalence of pathogenic variants. Several statistical methods exist to estimate MOI and the frequency distribution of pathogen variants. However, a common problem is the quality of the underlying molecular data. If molecular assays fail not randomly, it is likely to underestimate MOI and the prevalence of pathogen variants.

**Methods and findings:** A statistical model is introduced which explicitly addresses data quality, by assuming a probability by which a pathogen variant remains undetected in a molecular assay. This is different from the assumption of missing at random, for which a molecular assay either performs perfectly or fails completely. The method is applicable to a single molecular marker and allows to estimate allele-frequency spectra, the distribution of MOI, and the probability of variants to remain undetected (incomplete information). Based on the statistical model, expressions for the prevalence of pathogen variants are derived and differences between frequency and prevalence are discussed. The usual desirable asymptotic properties of the maximum-likelihood estimator (MLE) are established by rewriting the model into an exponential family. The MLE has promising finite sample properties in terms of bias and variance. The covariance matrix of the estimator is close to the Cramér-Rao lower bound (inverse Fisher information). Importantly, the estimator’s variance is larger than that of a similar method which disregards incomplete information, but its bias is smaller.

**Conclusions:** Although the model introduced here has convenient properties, in terms of the mean squared error it does not outperform a simple standard method that neglects missing information. Thus, the new method is recommendable only for data sets in which the molecular assays produced poor quality results. This will be particularly true if the model is extended to accommodate information from multiple molecular markers at the same time, and incomplete information at one or more markers leads to strong depletion of sample size.

## Introduction

The global COVID-19 pandemic underlined the importance of molecular surveillance in disease control and prevention, as reflected by the recently released WHO Global genomic surveillance strategy [1]. Molecular surveillance is well-established in some of the most relevant infectious diseases in terms of incidence, mortality, and economic burden. For instance, wide-scale malaria interventions led to major reductions in overall morbidity and mortality, a significant global decline in incidence during the past two decades, and a shift from disease control towards elimination. In this context, molecular surveillance to characterize transmission dynamics is becoming an essential tool. A popular quantity in molecular surveillance of malaria is multiplicity of infection (MOI). Namely, in this context, MOI or complexity of infection (COI) are established as a fundamental metric which scales with transmission intensity. Here, MOI is referred to as the total number of super-infections due to multiple infective contacts during one disease episode following the definition of [2] (see Figure 1). Note, the same pathogen ‘lineage’ might super-infect a host several times, which is accounted for by this definition. In malaria, the distribution of MOI is particularly important because it mediates the effective amount of recombination between haplotypes, and hence the population genetics of the parasite. Importantly, the concept of MOI is not restricted to malaria.

Several methods to estimate the distribution of MOI and frequency distribution of parasite lineages (e.g. allele or haplotype frequencies) from molecular data exist (see the introductions in [3] and [2] for an overview). In the case of malaria, molecular data is usually generated from disease-positive blood samples and contains allelic information at molecular markers. These markers are typically SNPs at specific positions, short SNP barcodes (micro-haplotypes), microsatellite markers, or var-gene expressions. Due to super-infections (i.e. MOI) several “alleles” can be observed at a molecular maker. The statistical properties of some of the existing methods to estimate MOI and lineage frequencies have been studied in detail (e.g. [3–5]). However, a common problem in practice is the quality of the molecular data itself – missing information at one or several molecular markers is common. Whether the performance of the statistical methods are affected by poor data quality depends on the nature of missing values.

The simplest case is to assume that for a given sample a molecular assay fails randomly to generate output. This implies missingness at random, in which case samples with missing information can simply be omitted in an analysis, leading to a reduction in sample size. However, there might be some more mechanistic reasons leading to incomplete information rooted in the molecular assays. Namely, each molecular assay has a detection threshold. Allelic variants which surpass this threshold are typically called, whereas those that stay below the threshold are considered noise. If in an assay allelic variants fail independently to amplify, missingness is no longer at random. In a sample the assay might capture the full, only partial, or no allelic information. It is obvious that an assay fails completely if no information is retrieved from a sample. However, if this is not the case, it is impossible to decide with certainty whether the full or only partial information was gained. In the following we use the term incomplete information, if no or only partial information is retrieved from a molecular assay. In any case, samples with no information can no longer be simply disregarded as missing at random from an analysis, and need to be adequately accounted for by a statistical method. Clearly, the specific mechanism jeopardizing data quality depends on various factors, e.g., including improper handling of clinical specimens, degradation of DNA over time, poorly performed assays, data entry errors etc. Incomplete information might be specific to the type of molecular data being generated, e.g., STR *vs*. SNP data. Knowledge about these factors can contribute to establishing a statistical model to account for incomplete information.

Here, in a first attempt we consider the simplest case of a single molecular marker to develop a statistical model to estimate MOI and lineage frequencies/prevalences from molecular data characterized by incomplete information. More precisely, we extend the maximum-likelihood method of [6] to account for incomplete molecular data. The model assumes that the lineages fail to be retrieved from a molecular assay independently of each other. Note, that the definition of MOI here accounts for multiple infectious events with the same ‘lineage’. Whether a lineage fails to amplify is assumed to be independent of how many times the lineage was super-infecting. These assumptions are justified because the proportions of different infecting pathogen lineages: (i) change due to intra-host dynamics; (ii) might not be correctly reflected in a clinical specimen; or (iii) change due to random events in the molecular assay (e.g. PCR reaction). The model estimates the distribution of MOI, assuming a conditional Poisson distribution, the frequencies and prevalences of the lineages, and the probability that a lineage remains undetected by a molecular assay.

In Methods the statistical model is formulated and the numerical investigations to evaluate the proposed methods are described. In Results it is shown that the statistical model belongs to the class of exponential families, by which we prove existence and uniqueness of the maximum-likelihood estimate except in pathological cases, which are characterized. This also proves the typical desirable asymptotic properties of maximum-likelihood estimators. There exists no explicit solution for the maximum-likelihood estimate, hence it needs to be calculated numerically. While this can be done straightforwardly by a multidimensional Newton-Raphson method, we advocate the use of the EM-algorithm here. Furthermore, the Fisher information is derived and it is described how to efficiently calculate the Cramér-Rao lower bound [7]. Finally, using numerical simulations, we report on the performance of the methods under finite sample size and compare it to the method of [6] under the-missing-at-random assumption.

## Methods

We consider a parasite population, whose pathogen variants are characterized by *n* different lineages (alleles) at a single marker locus or *n* distinct haplotypes in a non-recombining region. We refer to these pathogen variants as lineages which are circulating in the pathogen population. We consider only lineages that contribute to transmission, while we ignore those which are generated by mutation inside hosts and fail to participate in transmission. It is also assumed that the lineages are characterized by molecular markers, whose lineage-frequency distributions do not change too rapidly in the population, e.g., neutral molecular markers.

Let the *n* lineages be denoted by *A*_1_, …, *A*_*n*_, and their respective frequencies in the pathogen population by *p*_1_, …, *p*_*n*_. A host gets infected with exactly one lineage at each infective event. However, hosts can get super-infected by the same or different lineages. Let *m* be the number of super-infections referred to as multiplicity of infection (MOI). Hence, if *m*_*k*_ is the number of times that the host is infected by lineage *A*_*k*_, then nfection with MOI *m*, the pathogen configuration ***m*** = (*m*_1_, …, *m*_*n*_) corresponding *m* =*m*_1_+ … +*m*_*n*_. Furthermore, we assume that at each infective event a lineage is drawn randomly and independently from the pathogen population. Hence, for an infection with MOI *m*, the pathogen configuration ***m*** = (*m*_1_, …,*m*_n_) corresponding follows a multinomial distribution with parameters *m* and ***p***, where ***p***= (*p*_1_, …, *p*_*n*_) is the vector of lineage frequencies. If infective events are rare and independent, it is shown by a queuing model that MOI follows a Poisson distribution [8]. Clearly, no infection (or *m*= 0) corresponds to a disease-free individual. Hence, among disease-positive individuals MOI follows a conditional Poisson distribution (CPD), i.e.,

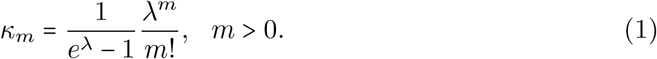

This distribution is characterized by the Poisson parameter *λ*. The average MOI is derived as

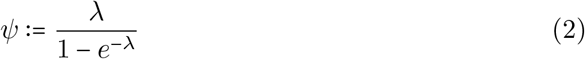

(cf. [9]).

### Ideal observations

In practice, it is impossible to observe the number of times an individual is infected by each lineage, i.e., the values corresponding to *m*_*k*_ cannot be observed in an infection (see Fig 1 “unobservable information”). However, the absence/presence of lineages can be observed in a clinical sample. Let *y*_*k*_ ∈{0, 1} denote the absence/presence of lineage *A*_*k*_ in a blood sample. Therefore, an observation (sample) represented by the configuration

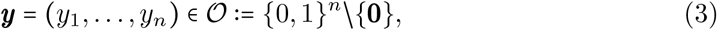

where **0** is the *n*-dimensional vector with all zero components, and represents a clinical sample in which no lineage is detected (this configuration is excluded, because we only consider disease-positive individuals). The probability of a sample having configuration ***y*** is given by

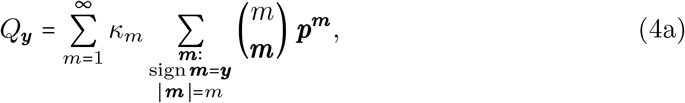

or

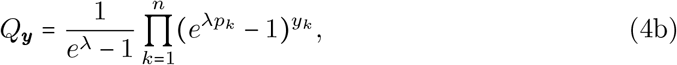

(cf. [9]). This model is identifiable, i.e., different sets of parameters lead to different distributions of ***y*** (cf. [4]). Moreover, observations are “ideal” because we assume that all lineages actually present in an infection will be detectable and lineages are not erroneously detected.

**Fig 1.**
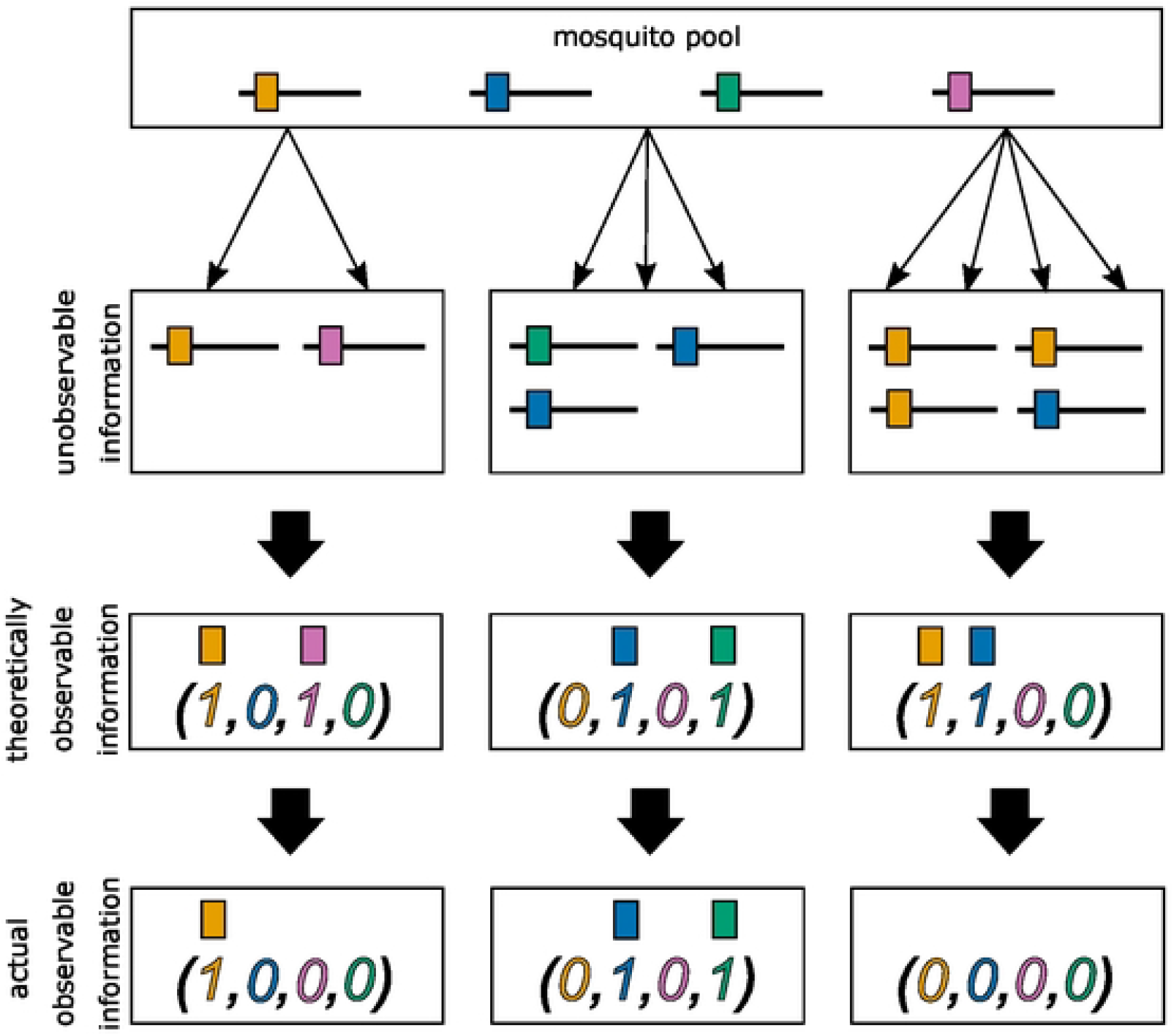
Illustration of incomplete data. Illustration of information contained in blood samples. The top row illustrates three (super-)infections. The middle row illustrates the respective information that is theoretically observable from a blood sample. The bottom row illustrates hypothetical cases for which miss-observation can occur. The first individual (left column) was infected by *m* =2 lineages, one time with the orange and once with the pink lineage. These are in principle observable in a blood sample. However, the molecular assay detected only the orange lineage, while the pink one remains undetected. In the middle, a super-infection with *m* =3 lineages is illustrated. In this case no lineage remains undetected and all lineages present in the infection are observed. The third individual is infected with orange (three times) and blue (one time) lineages. However, the actual observation shows that the individual is not infected at all. This is the case of complete missing information.

### Incomplete observations

In practice, a lineage within an infection might remain undetected. To accommodate this important factor, we extend the model. We assume that lineages remain undetected independently of each other and detection of each lineage is independent of the true number of super-infections by that lineage, *m*_*k*_. We assume further that each lineage is equally likely to be detected. Hence, we let *ε* be the probability that a lineage, present in the infection, remains undetected in the blood sample. Consequently, the probability of a lineage being detected correctly is 1 −*ε*. Additionally, we ignore that lineages which are absent from an infection can be detected by mistake. Under these assumptions, the true absence/presence of lineages ***y*** remains unobserved, and the corresponding actual observation (denoted ***x***) might not correctly indicate the presence of all lineages being actually present, i.e.,

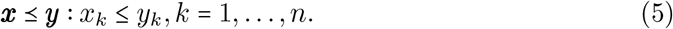

Therefore, for an observation ***x***, it remains unclear which ***y*** reflects the true absence/presence of lineages (see Fig 1 “actual observable information”). Thus, the probability of observing ***x*** has to take into account that any ***y*** with ***x*** ⪯ ***y*** can lead to detected then ***x*** = 0. We refer to **0** as the empty record. According to above observing ***x*** with lineage *A*_*k*_ missing if *y*_*k*_ = 1 but *x*_*k*_ = 0. If all lineages fail to be assumptions, given that ***y*** reflects the true absence/presence of lineages in an infection, the probability of observing ***x*** is

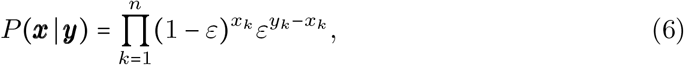

for ***x***⪯ ***y*** and zero otherwise. As for every observation ***x*** each ***x*** ⪯***y*** potentially reflects the true absence/presence of lineages, according to the law of total probability, the probability of observing configuration ***x*** =(*x*_1_, …, *x*_*n*_) thus becomes

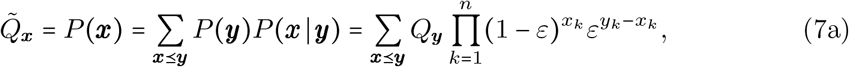

where *Q*_***y***_ is specified in (4b). The sum runs over all possible ideal observations that can lead to to the actual observations, i.e., for all ***x*** ⪯***y*** (i.e., *x*_*k*_≤ *y*_*k*_ for all *k*= 1, …, *n*). It is shown in section Prevalence in S1 Mathematical appendix that 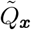 can be expressed as

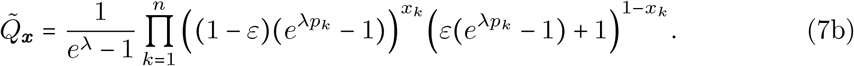

By accounting for incomplete data, the empty record **0** does not correspond to a disease-free host, as only samples from infected hosts are considered. The empty record occurs with probability

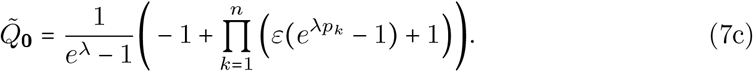

We refer to the new model (7) as the incomplete-data model (IDM). It includes the additional parameter *ε* compared with the original model (OM). The sample space of the IDM is denoted by 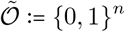.

Jointly we denote the model parameters by ***θ*** = (*λ, p, ε*). Clearly, the MOI parameter *λ* is strictly positive. The frequency vector ***p*** satisfies 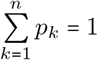 and 0 < *p*_*k*_ < 1 for *k* =1, …, *n*, and therefore is an element of the interior of the (*n* − 1)-dimensional simplex. Additionally, we assume *ε*∈(0, 1). Hence, the parameter space of the new model becomes

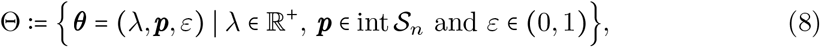

where int **𝒮**_*n*_ denotes the interior of the (*n* −1)-dimensional simplex. Note that, if *ε* = 0, no lineage remains undetected and all lineages present in an infection will be observed almost surely. In this case, the IDM coincides with the OM. Additionally, if *ε* = 1, (6) implies that 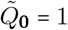 and 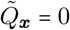 if 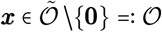 for any choice of *λ* and ***p***, and hence, the model is not identifiable.

### Prevalence

It is important to point out that *p*_*k*_ is the relative frequency of lineage *A*_*k*_ and not its prevalence. The former refers to the relative abundance of the lineage in the pathogen population, while the latter refers to the probability of the lineage’s presence in an infection (within the population of disease-positive individuals). The (actual) prevalence of lineage *A*_*k*_, i.e., the probability that the lineage occurs in a samples, irrespective of incomplete observations, is

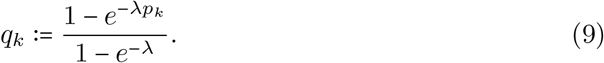

Accounting for incomplete data, the observable prevalence is not just mediated by MOI, but also by the probability of lineages not being detected (*ε*).

#### Remark 1.

*Under the IDM, the observable prevalence of lineage A*_*k*_, *i*.*e*., *the probability of observing A*_*k*_ *in a blood sample, is*

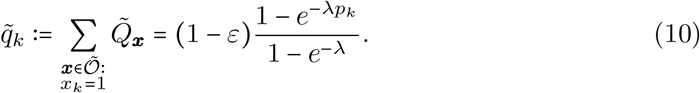

*The observable prevalence accounts for the possibility that a lineage remains unobserved because of incomplete information*.

*The probability of observing lineage A*_*k*_ *in samples, excluding empty records, i*.*e*., *the observable prevalence condition on not observing empty records (referred to as conditional prevalence), is given by*

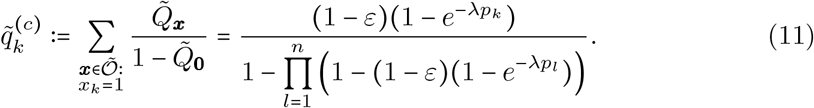

A proof is provided in section Prevalence in S1 Mathematical appendix. If super-infections occur rarely, the prevalence of lineages are expected to be close to their corresponding frequencies. In fact, in the limit as *λ*→ 0 these quantities coincide (see section Prevalence in S1 Mathematical appendix).

### The likelihood function

Let **𝒳** be a dataset containing the information obtained from *N* samples. That is, **𝒳** consists of *N* 0-1 vectors, where each vector corresponds to an observation indicating the detected lineages. Let 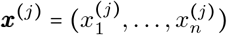 denote the observed data corresponding configuration ***x*** in the dataset **𝒳**. Thus, if ***x***^(*j*)^ ≠ ***x*** for *j* = 1, …, *N*, we have *n*_***x***_ = 0. In to the *j*-th sample. Furthermore, let *n*_***x***_ denote the number of samples with particular, *n*_**0**_ is the number of empty records in **𝒳**. Clearly

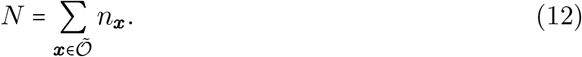

The log-likelihood function for the observed dataset X is given by

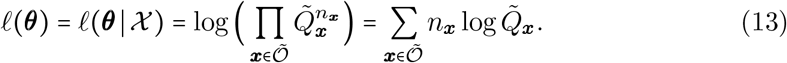

Let

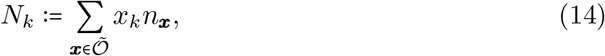

be the number of samples for which lineage *A*_*k*_ is observed. It is shown in section Log-likelihood function in S1 Mathematical appendix that the log-likelihood function (13) can be simplified to

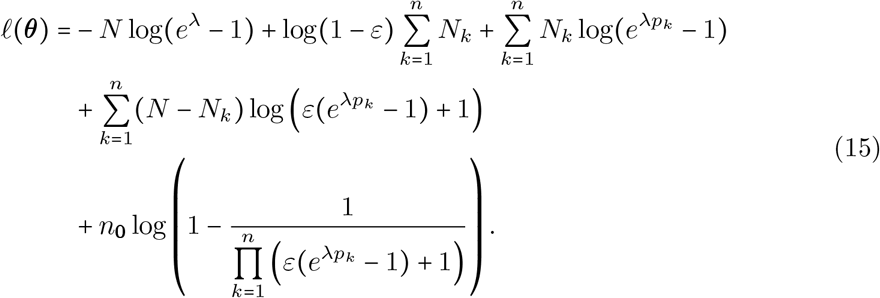

The sample size *N* can be written as *N* = *n*_**0**_ + *N*_+_, where

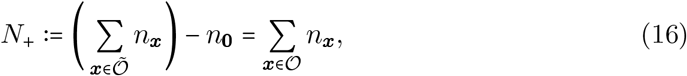

is the number of samples for which at least one lineage is observed – or the number of non-empty records. Clearly, the quantities *n*_**0**_, *N*_1_, …, *N*_*n*_ form a sufficient statistic for the IDM. More importantly, 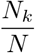 is the (unconditional) observable prevalence, and 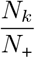 is the conditional observable prevalence of lineage *A*_*k*_ in the pathogen population.

### Assessing the performance of the model extension

The advantage of the extended model (7) in comparison to the OM (4) is that it allows to incorporate empty records, i.e., all samples in a dataset can be included in the parameter estimation. When parameters are estimated from the OM, empty records have to be excluded from the data, because such observations are not accounted in the model. The advantage of the model extension (7) is assessed here numerically.

Numerical investigations are conducted for a representative range of model parameters (summarized in Table 1) as follows. For a set of parameters (*N, λ*, ***p***, *ε*) a total number of *B* = 100 000 regular datasets (cf. Pathological Cases) **𝒳**_1_, …, **𝒳**_*B*_ of size *N* are created according to the model (7). Each dataset **𝒳**_*b*_ might contain empty records. Let 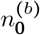 and 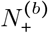 be the number of empty and non-empty records of **𝒳**_*b*_. Let further 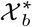 be the dataset **𝒳**_*b*_ with all empty records removed. Thus, the sample size of 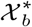 is 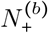. For each dataset **𝒳**_*b*_, the maximum-likelihood estimate (MLE), 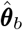, based on the extended model (7), is calculated (see Deriving the maximum-likelihood estimate). Furthermore, from the MLE 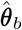 the average MOI 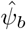 is calculated according to (2).

**Table 1.** Summary of model parameters. Shown are the parameter values used to generate simulated datasets. A total number of *B* datasets were generated for each combination of model parameters and sample size (*N*).

| Summary of Model Parameters for the Simulation Study |  |  |
| --- | --- | --- |
| par. | description | values |
| $B$ | number of generated datasets | 5 000 |
| $\lambda$ | Poisson parameter | 0.1, ..., 2.5 in steps 0.1 |
| $\psi_{CP}$ | average MOI | $\frac{\lambda}{1 - e^{-\lambda}}$ with $\lambda$ as above |
| $N$ | sample size | 50, 100, 150, 200 |
| $n$ | number of lineages | 2, 3, 4, 5 |
| $\varepsilon$ | prob. that lineages remain undetected | 0.015, 0.10, 0.15 |
| $\mathbf{p}$ | lineage-frequency distributions: | |
| | $n = 2$ | (0.5, 0.5), (0.75, 0.25) |
| | $n = 3$ | (1/3, 1/3, 1/3), (0.75, 0.15, 0.1) |
| | $n = 4$ | (0.25, 0.25, 0.25, 0.25), (0.75, 0.15, 0.05, 0.05) |
| | $n = 5$ | (0.2, 0.2, 0.2, 0.2, 0.2), (0.75, 0.1, 0.05, 0.05, 0.05) |

Furthermore, for each dataset 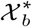 the MLE 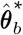 of the parameter vector ***θ***^***\****^ is calculated based on the methods described in [9], i.e., based on the OM. From 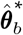 the average MOI 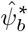 was calculated according to (2).

Bias and variance of the estimators for both models (IDM and OM) were estimated as follows. Denoting a model parameter generically by *θ* (i.e., *θ* is *λ, ψ, p*_*k*_, or *ε*), the bias is estimated as

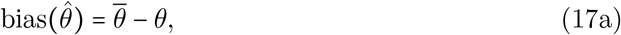

where

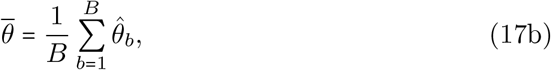

and *θ*_*b*_ is the corresponding MLE from the *b*-th dataset. The variance of the the estimator for *θ* is calculated as

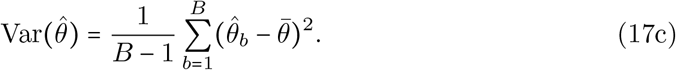

Because bias and variance of the estimators depend on the range of the true parameters, the relative bias and coefficient of variation, i.e.,

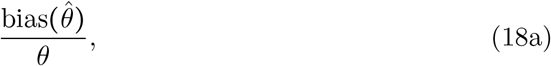

and

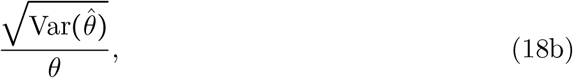

are more appropriate for comparison of the bias and variance across different parameter ranges.

Additionally, we derived the empirical mean-squared error to measure the overall quality of the estimators. Similarly, to be able to compare the approximated mean-squared error of an estimator across a range of *ψ* values, we used the normalized root-mean-square error (NRMSE), i.e.,

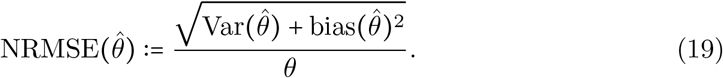

Clearly, (i) sample size *N*, (ii) the true value of the MOI parameter *λ*, (iii) the number of lineages *n*, (iv) the lineage-frequency distribution ***p***, and (v) the probability that a lineage remains undetected *ε* affect bias and variance of the MLEs. For the numerical investigations, we choose the following parameters.

#### Sample size

The main difference between the two models is that the extended model incorporates empty records. The sample size of a simulated dataset is reduced in the presence of empty records when using the OM. We consider sample sizes *N*= 50, 100, 150, and 200 for the extended model.

#### MOI parameter

Concerning the MOI parameter we chose *λ*= 0.1, …, 2.5 with increments of 0.1, which corresponds to an average MOI *ψ* ranging from 1.05 to 2.72. In the case of malaria, low transmission corresponds to *ψ* < 1.27 (*λ* = 0.5), intermediate transmission to 1.27 ≤ *ψ* < 1.93 (0.5 < *λ* ≤ 1.5), an high transmission to *ψ* ≥ 1.93 [3].

#### Number of lineages

The number of lineages also contributes to an estimator’s performance. As *n* increases, the likelihood of observing empty records decreases, and the two estimators converge in performance. Based on these facts, we considered lineage-frequency distributions with *n*= 2, 3, 4, and 5.

#### Lineage-frequency distribution

It is also important to study the performance of the MLE with respect to the underlying lineage-frequency distribution. Notably, cases for which lineages are similar in frequency as opposed to the cases for which one lineage predominates, need to be explored. For the former we consider a completely uniform (balanced) distribution, i.e., all lineages have the same frequency,

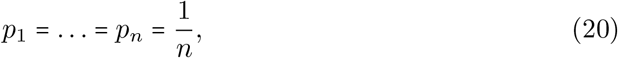

whereas for the latter an unbalanced distribution with one predominant lineage is considered. The frequency of the predominant lineage is chosen to be 75%, i.e., *p*_1_ =0.75. The distributions are summarized in Table 1.

#### Probability that a lineage remains undetected

The probability that a lineage remains undetected is chosen to be *ε* =0.05, 0.01, and 0.15.

Because we only generated regular datasets, all inferences have to be understood as conditioned on regular data.

We used R [10] to implement the simulation study and create the graphical outputs. The code necessary to reconstruct the simulations is available at https://github.com/Maths-against-Malaria/MOI---Incomplete-Data-Model.git.

## Results

### Maximum likelihood estimate and Fisher information

Here we discuss approaches to obtain the maximum-likelihood estimate (MLE) for the model parameters ***θ*** = (*λ, p, ε*). The first approach is to maximize the likelihood function (15) directly. Since the frequencies ***p*** are an element of the simplex **𝒮**_*n*_, the log-likelihood function (15) needs to be maximized over the parameter space Θ, either by using a Lagrange multiplier or by eliminating one of the redundant variables, e.g., by setting 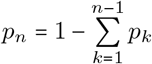. Using a Lagrange multiplier, the log-likelihood function (15) is replaced by

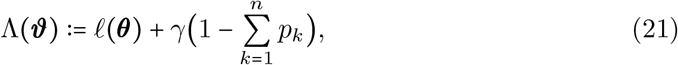

where ***ϑ*** = ***(θ***, *γ)*. The MLEs of the model parameters are found by equating the score function to zero, i.e., by solving the following system of equations

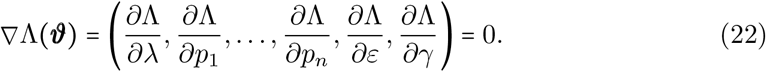

Unfortunately, the system ∇Λ (***ϑ)* = 0** has no explicit solutions and needs to be solved recursively, e.g., by a multi-dimensional Newton-Raphson method, i.e., by iterating

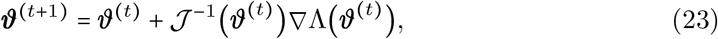

where **𝒥*(ϑ***^(*t*)^) is the observed information (negative of the Hessian matrix of Λ (***ϑ)***) evaluated at ***ϑ***^(*t*)^. The score function and the observed information **𝒥** are given in section Inverse Fisher information in S1 Mathematical appendix.

As seen in section Inverse Fisher information in S1 Mathematical appendix, the data (*N*_1_, …, *N*_*n*_, *n*_**0**_) (which are random variables) occurs only in linear terms in the observed information. Because the Fisher information is the expected observed information, it can be easily calculated because 𝔼*N*_*k*_ and 𝔼*n*_**0**_ follow readily. Namely, 𝔼*N*_*k*_ is the expected number of infection in which lineage *k* is present. Because the probability to observe lineage *k* in a sample is its prevalence 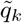, *N*_*k*_ is a binomially distributed random variable with parameters *N* and 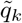. Similarly, *n*_**0**_ is binomially distributed with parameters *N* and 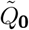. Therefore,

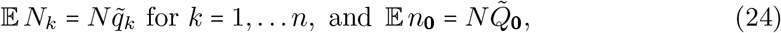

where 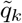 and *Q*_**0**_ are given by (10) and (7c), respectively. The resulting entries of the Fisher information are presented in section Inverse Fisher information in S1 Mathematical appendix.

Both the Fisher information and the observed information contain the Lagrange parameter *γ* as a nuisance parameter. The reason is that the structure of the matrix is simpler. Namely, it contains a diagonal matrix corresponding to the second derivatives with respect to the lineage frequencies as a sub-matrix. This structure allows straightforward inversion by using block-wise inversion formula [11]. The Cramér-Rao lower bound (cf. [7]) is the inverse Fisher information matrix (with the entries corresponding to the nuisance parameter disregarded) and yields the covariance matrix of the estimator. The Cramér-Rao lower bound yields the minimum variance of any unbiased estimator, and will be asymptotically reached by the MLE. Although it is straightforward to invert the information matrix, the resulting formulae are lengthy. Consequently, we refrain from presenting analytical expressions here and refer S1 Mathematical appendix for the expressions of the information matrices.

In practice, the true parameter is unknown, so either the inverse Fisher information or the inverse observed information have to be estimated using the MLE as a plug-in estimate. It follows from Remark 2 and the derivations in section Inverse Fisher information in S1 Mathematical appendix that the estimates for the inverse Fisher information and the inverse observed information coincide. One might be more interested in the average MOI *ψ* (cf. eq. 2) rather than in the MOI parameter *λ*. To obtain the covariance matrix with respect to this estimator the inverse Fisher information needs a slight adjustment (see Inverse Fisher information in S1 Mathematical appendix).

We can conclude that it is advisable to use the method of Lagrange multipliers to derive the covariance matrix of the estimator. To derive the estimate itself, the Newton method (23) is not advisable. Even it is replaced by a quasi-Newton method or other modified versions of the Newton method, convergence of these methods is too sensitive on initial conditions. For such systems, usually sufficiently accurate initial values are required for convergence.

We will pursue with the expectation-maximization (EM) algorithm. The EM algorithm turns out to be convenient in our case. However, the EM algorithm has a tendency of approaching the maximum fast, but convergence might slow down close to the maximum. As an improvement a combination of the EM algorithm and a Newton method can be used (cf. [12] chapter 2).

Before doing so we will rewrite the model into an exponential family. We will show that the model belongs to a regular and natural exponential family, which concludes that the EM algorithm finds the unique MLE for the model parameters. Additionally, rewriting the model as an exponential family helps us to understand the restrictions of the model, i.e., the cases for which the MLE does not exist.

### Model as a natural exponential family

The model (7a) can be rewritten as a natural exponential family [13]. Consequently, the log-likelihood (15) also can be rewritten in this form, since the exponential families are preserved under repeated sampling, given that the observations are independent.

Remember, for the IDM we defined the sample space of all possible observations as 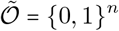. While 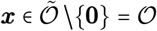 retains some allelic information about the underlying data, even if the information is incomplete, the empty record **0** harbors no ‘direct’ information. This particular observation deserves special attention. We transform the sample space 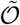 into a higher-dimensional space for which the first component specifies whether a record is empty. More precisely, a non-empty observation ***x*** ∈ **𝒪** is identified with the vector (0, ***x***) and the empty record with the vector (1, **0**). Let

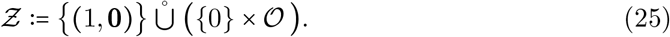

We define the transformation 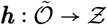 by

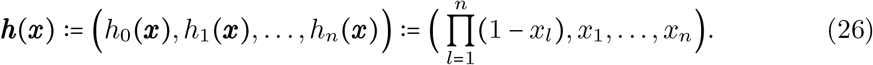

It is easy to see that ***h***(***x***) is a bijection between 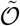 and **Ƶ**. Note, *h*_0_(***x***) serves as an indicator for the empty record, i.e., *h*_0_(**0**) = 1 and *h*_0_(***x***) = 0 for all ***x*** ∈ **𝒪**.

Let the vector-valued function ***g* :** Θ → Ω, where 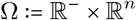, be defined as

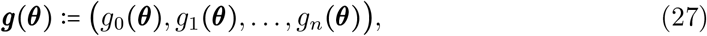

with

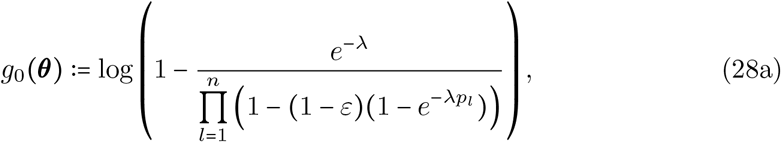

and

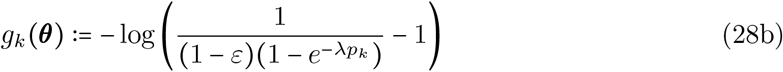

for *k* = 1, …, *n*. Additionally, let

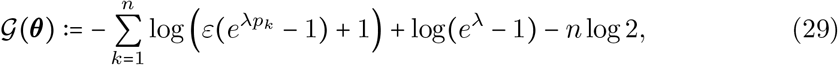

and

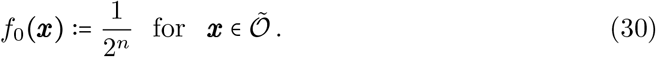

It is shown in section Incomplete-data model (IDM) as a natural exponential family in S1 Mathematical appendix that the model (7a) can be rewritten as an exponential family in the form

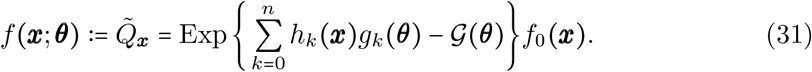

The function *f*_0_ (***x)*** is the base density, which is a uniform density on the support of ***x***. This representation is not in natural form. It is possible to rewrite (31) into the natural form by substituting ***β*:**= ***(g θ)*** since ***g*** is a 1-1 mapping (see section Incomplete-data model (IDM) as a natural exponential family in S1 Mathematical appendix).

Let ***z***^*T*^ ***β*** denote the scalar product of ***z*** and ***β***, i.e.,

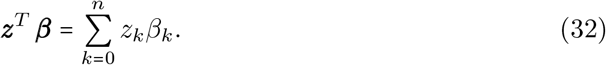

Hence, the model (31) is rewritten into its natural form as

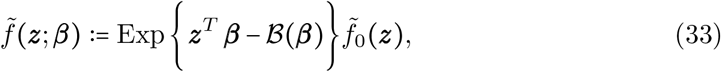

where

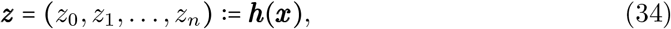

***β*** = (*β*_0_, *β*_1_,…, *β*_*n*_) with β_k_ = *gk*(**θ**) for *k* = 0, 1,…, *n*,

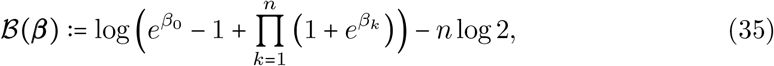

and 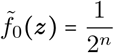 is the base density of the natural exponential family (see section Incomplete-data model (IDM) as a natural exponential family in S1 Mathematical appendix).

Because ***h (x)*** is a bijection, the marginal distribution of ***z*** is equivalent to the distribution of ***x***, i.e., both have the same structure function (base density). The closed convex hull of its support is

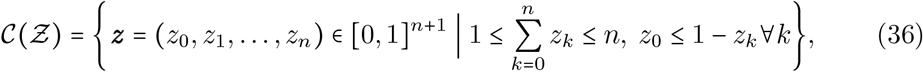

where **Ƶ** is specified in (25). It is not difficult to see that ***h*** is also bijection between **𝒞** (**Ƶ)** and the closed convex hull of 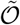, which is an *n*-dimensional rectangular set in ℝ^*n*^.

The function **ℬ**(***β***) can be interpreted as the cumulant-generating function of ***z*** (cf. [7], chapter 5.2.1). The natural parameter space of the exponential family is the set for which **ℬ*(β)*** is finite. From (35) it can be seen that this is ℝ^*n*+1^. This implies that the natural exponential family (33) is regular. Additionally, the natural statistic ***z* = *h (x)*** and the natural parameter space are (*n+*1) -dimensional, showing that the natural representation is in fact minimal (see section Minimality of the natural exponential family in S1 Mathematical appendix). The minimality of (33) ensures that the natural exponential family is full, not curved, and that the statistic ***z*** is minimal sufficient for ***β*** [13].

Now, consider a dataset **𝒳** consisting of *N* observations ***x***^(1)^, …, ***x***^(*N*)^ obtained from *N* disease-positive samples, such that each ***x***^(*j*)^ (for *j* = 1, …, *N*) is distributed according to the exponential family (31) with the common corresponding parameter space Θ. The corresponding natural statistics are the observations ***z***^(1)^, …, ***z***^(*N*)^ with ***z***^(*j*)^ ***h (x***^(*j*)^), which follow the natural exponential family (33) with natural parameter space ℝ^*n*+1^. The likelihood of observing **𝒳** as a function of ***β*** is

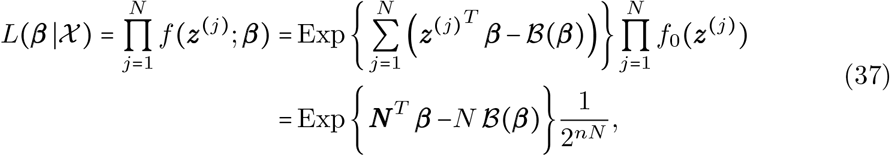

where

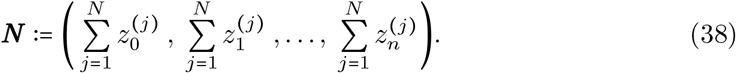

Note that 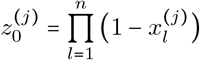 is the indicator of the *j*-th sample being an empty record, and for 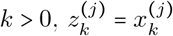 indicates if lineage *A*_*k*_ was observed in sample *j*. Hence,

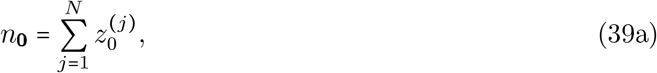

and

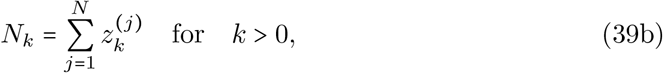

where *n*_**0**_ is the number of empty records, and *N*_*k*_ is the number of samples infected by lineage *A*_*k*_ (one can easily conclude that this is an equivalent representation for eq. 14).

Therefore, (38) becomes

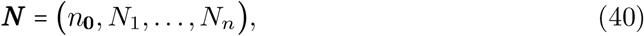

which is a sufficient statistic for the likelihood (37). In fact, ***N*** is minimal sufficient, since the natural exponential family (33) is minimal [13]. Clearly, the likelihood has also the natural exponential family of the form (33).

Let

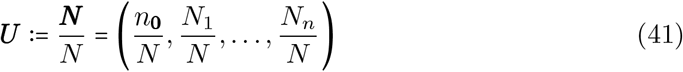

be the vector of empirical prevalences. The log-likelihood function is derived as

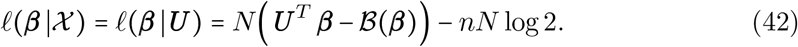

It is easy to see that the support of ***U*** is

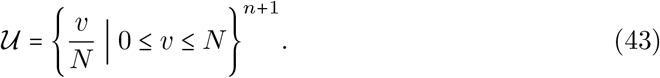

The cumulant-generating function of ***U*** is the same as the cumulant-generating function of ***z***. Because the family is regular, it is strictly convex and sufficiently many times differentiable [13]. Its first derivative with respect to the natural parameters is the mean vector of ***U***, i.e.,

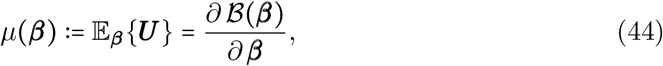

(cf. [14]), where the partial derivatives are

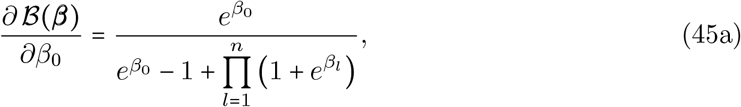

and

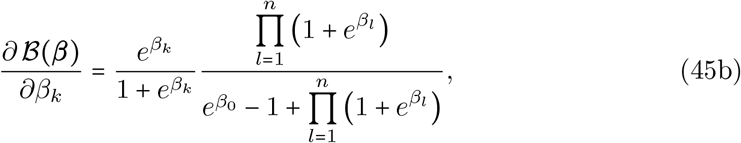

for *k* = 1, …, *n*. The log-likelihood function (42) is a strictly concave function of ***θ*** (it differs from − *B*(***β***) only by a linear term). Its first derivative (or the score function) is derived as

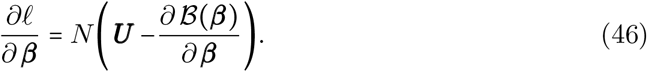

The MLE of 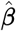 is then found as the unique root of

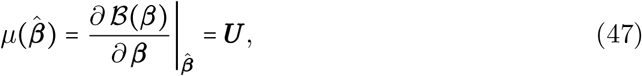

i.e., 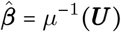. Therefore, 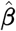 is derived by solving

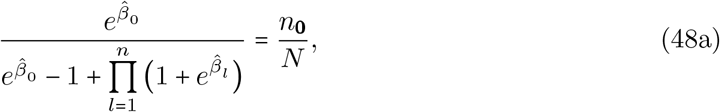

and

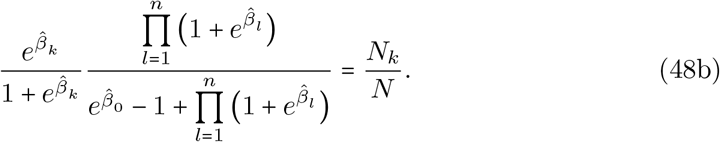

Since the natural model is regular, 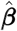 exists if ***U*** belongs to the interior of the closed convex hull of the support of ***U*** [15], which is the set

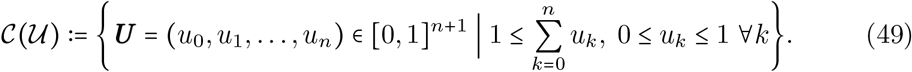

We will treat all pathological cases in which the MLE does not exist in the admissible parameter space in the next section.

In general the MLE yields a simple and intuitive interpretation, which is summarized in the following result.

#### Result 1.

*For the IDM* (4b), *the MLE is the choice of parameters* 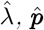, *and* 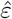, *for which (i) the probability of observing an empty record equals the observed relative frequency of the empty record, i*.*e*.,

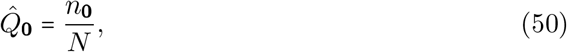

*(ii) the conditional prevalence of lineage k equals its relative frequency among non-empty records*,

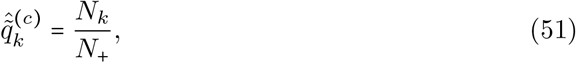

*and (iii) the relative frequency of samples containing lineage k among all (empty and non-empty records) is the actual prevalence multiplied with the probability that a lineage being present in an infection is actually observed, which equals the observed prevalence, i*.*e*.,

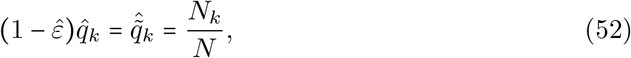

*where N*_*k*_, *n*_**0**_, *and N*_+_ = *N − n*_**0**_ *are considered as realizations of the corresponding random variables, i*.*e*., *as constants*.

The proof is presented in section Interpretation of the MLE in S1 Mathematical appendix.

#### Remark 2.

*The data N*_*k*_, *n*_**0**_, *and N*_+_ = *N* − *n*_**0**_ *is a realization of the random variables* 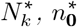, *and* 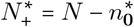. *From Result 1 and Remark 1 and* (24) *it follows that*

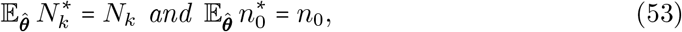

*where the expectation is taken with respect to the MLE* 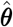. *In other words, of the MLE was the true parameter the expected number of samples containing lineage k equals the observed number and the expected number of empty records equals the observed number*.

*As a consequence the estimates for the inverse Fisher information and the inverse observed information based on the MLE coincide (cf. section Maximum likelihood estimate and Fisher information and section Inverse Fisher information in S1 Mathematical appendix)*.

### Pathological Cases

The MLE 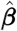 exists and is unique if ***U*** ∈ int **𝒞**(**𝒰)**. Importantly, of primary interest is 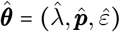, the MLE of the original parameters. We must be more cautious in deriving the MLE of ***θ***, since it only exists if 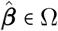. However, for some ***U*** ∈ int**𝒞**(**𝒰)** it is possible that 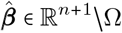. Therefore, the set of ***U*** for which 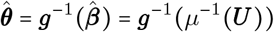 exists is only a subset of int **𝒞**(**𝒰)** (interior of the closed convex hull of **𝒰**). On the contrary, if ***U*** ∉int **𝒞**(**𝒰)**, 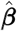 does not exits. However, these cases are interpreted in a limit sense. We refer to data ***U*** for which 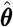 does not exits as pathological data. These cases are fully classified in Result 2.

First, it is worthwhile mentioning that if *N*_*k*_ = 0 for one *k* the model can be reduced by eliminating the parameter *p*_*k*_. Thus, we restrict our attention to *N*_*k*_ > 0 for all *k*.

#### Result 2.

*The following pathological cases occur*.

*Case A. Assume at least one lineage is observed in every sample, i*.*e*., *n*_**0**_ = 0. *The MLE coincides with those of the OM, which ignores incomplete data. The following four pathological cases occur*.

*A*.*1. If at least one sample contains evidence of a multiple infection, but no lineage is observed in every sample, in technical terms* 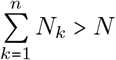 *and* 0 < *N*_*k*_ <*N for all k, then the MLE coincides with that of the OM. More precisely*, 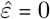 *and the estimates* 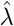 *and* 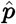 *are obtained as in [9]*.

*A*.*2. If there is no indication of multiple infections in the data, and no lineage is present in all samples, i*.*e*., *if* 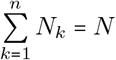 *and* 0< *N*_*k*_< *N for all k, the MLE is attained at*

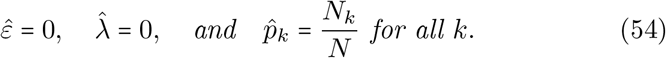

*Hence, the MOI parameter is estimated to be zero, and the lineage frequencies by their relative abundance in the data*.

*A*.*3. If at least one lineage occurs in all samples (and at least two lineages are observed in the data), i*.*e*., *if N*_*k*_ = *N and N*_*j*_ > 0 *for at least one j* ≠ *k, then* 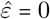 *and the MLE of the remaining parameters are reached in the limit λ* → ∞, *such that*

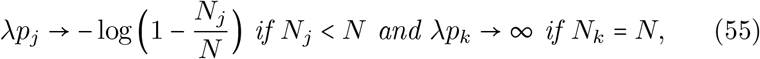

*subject to the constraint* 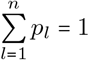.

*A.4*. *If only one lineage is observed in the data, i*.*e*., *N*_*k*_ *≠*0 *for only one k, the MLE is not unique and realized for any point*

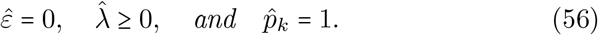

*Case B. Assume the fraction of empty observations is not “too large”, more specifically*, 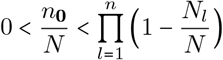. *The only pathological case arises under this condition if there is no indication of multiple infections in the data, and no lineage is present in all samples. More precisely, if* 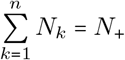 *and* 0 <*N*_*k*_ <*N*_+_ *for all k, the MLE is attained at*

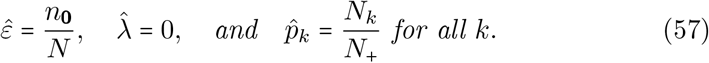

*Hence, the probability that a lineage remains undetected is estimated as the fraction of empty records, the MOI parameter as zero, and the lineage frequencies by their relative abundance in the non-empty observations*.

*Case C. If the fraction of empty records is relatively large, specifically*, 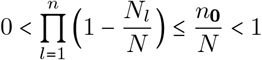 *the MLE is obtained in the limit case λ* → ∞*such that*

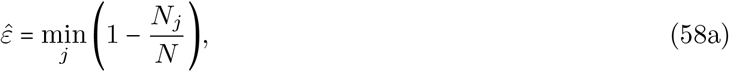

*and*

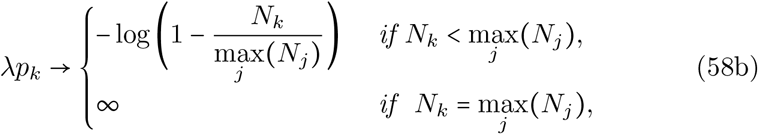

*subject to the constraint* 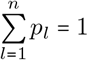. *In this case the supreme of the likelihood function is*

*For sake of completeness, in the case where all samples failed to produce data, i*.*e*., *n*_**0**_ **=** *N, the MLE is not well defined. In particular any point*

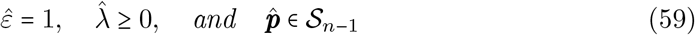

A proof of the results can be found in section Pathological cases in S1 Mathematical appendix.

### Deriving the maximum-likelihood estimate

By rewriting the statistical model into a natural exponential family, we could classify all pathological cases in which the MLE does not exist. Moreover, it has enabled us to establish Result 1, which gives an intuitive interpretation of the MLE in terms of prevalence but not in terms of the actual parameters of interest namely the ***θ* = (***λ, p, ε*). These need to be derived numerically because (44) has no explicit solution. However, the exponential family form yields a much simpler algorithm to derive the MLE than the multi-dimensional Newton-Raphson method (23), which is too sensitive to initial conditions. In terms of the exponential family, two 1-dimensional Newton algorithms need to be iterated. However, the problem of sensitivity to initial conditions is not entirely remediated. The EM-algorithm is an alternative, with the convenient property that convergence to the MLE is guaranteed except in the pathological cases. It is described in the following result.

#### Result 3.

*Given regular data, the MLE of the model parameters is obtained by a two-step iterative procedure. Starting from initial values λ*^(0)^, 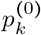 *and ε*^(0)^, *in step t* + 1 *the parameters p*_*k*_ *and ε are updated by*

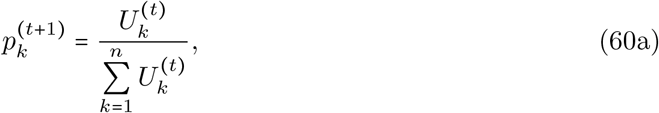

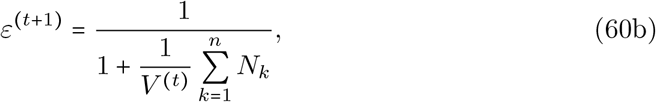

*where*

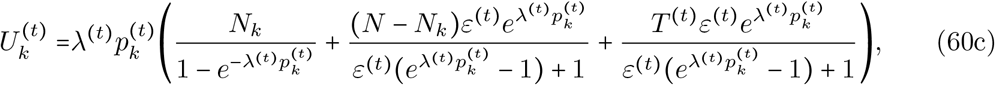

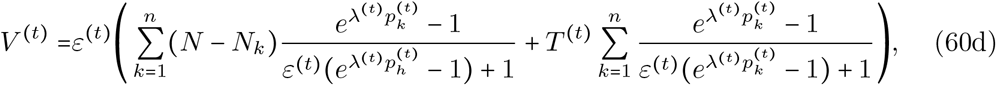

*and*

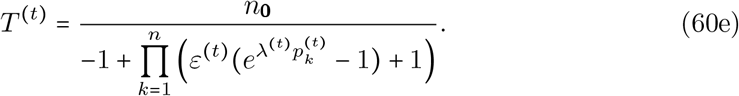

*The parameter λ is updated in step t* 1 *by iterating the recursion*

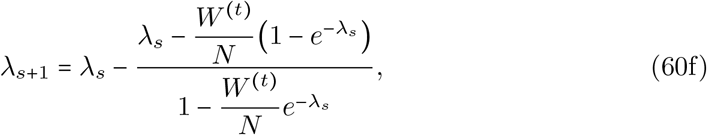

*starting from λ*_0_ = *λ*^(*t*)^, *where*

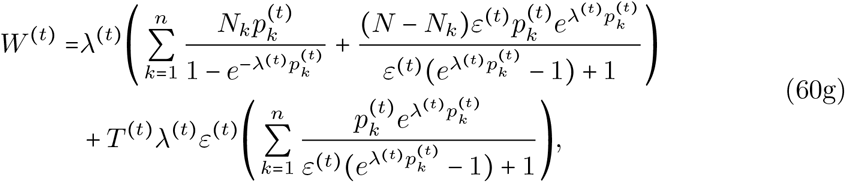

*until convergence is reached. This is the case if* |*λ*_*s*+1_ −*λ*_*s*_| <*δ, where δ is the numerical threshold for convergence. Then the parameter lambda is updated by λ*^(*t*+1)^ = *λ*_*s*+1_.

*The EM algorithm is guaranteed to converge monotonically from any initial values* ***θ***^(0)^ = (*λ*^(0)^, ***p***^(0)^, *ε*^(0)^). *Numerical convergence is reached according to a specified criteria, e*.*g*.,

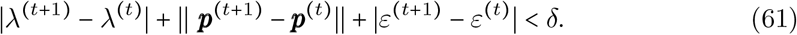

A complete outline of the calculations is given in section The EM Algorithm in S1 Mathematical appendix.

Note that the EM algorithm needs 1-dimensional Newton algorithm in the maximization step. This Newton algorithm finds a unique solution in every iteration if the data is regular. The MLE derived from the EM algorithm exists and it is unique, since the log-likelihood function belongs to a natural and regular exponential family [16]. However, the EM algorithm does not converge (i.e., the MLE does not exist) in the pathological cases.

### Finite sample properties

Concerning the finite sample size properties of the proposed method, we first concentrate on bias. To be able to compare the bias of the MOI parameter *λ* or rather the average MOI *ψ* across different parameter ranges, the relative bias (i.e., the bias in percent of the true underlying parameters) is reported. Regarding the lineage frequencies we report the bias of the predominant lineage. Second, the variance of the estimators, in terms of the coefficient of variation are reported, which again allows a comparison of the performance between different parameter ranges. The performance of the IDM is also compared to that of the OM.

Note that the bias and variance of the OM, were already studied in [3, 4]. However, it was studied under the assumption of complete data, i.e., *ε* =0. Once *ε*> 0, the OM is no longer the true model. Namely, the quality of observations is reduced, because of missing information, i.e., not all lineages in a sample are observable. As shown in Remark 1 if no empty records occur in the dataset, the IDM and the OM yield the same estimates, since they use the same data. In particular, the IDM wrongly estimates *ε* as 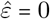. If empty records occur, the IDM and the OM yield different estimates and the OM is based on sample size *N*_+_ rather than on *N*.

#### Bias of the average MOI

In the case of perfect information *ε =* 0 the IDM and OM coincide, and the bias was described already in [3, 4], for which the OM and IDM coincide.

The bias of the average MOI (Figs 2, 3) estimated by the IDM is rather insensitive to the probability of lineages remaining undetected, i.e., to *ε*, and remains similar as described in [3, 4]. More precisely, if *ε* increases from 1.5% to 15%, the maximum bias increases only by 3%. The bias of 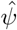 is a non-linear function of the true average MOI *ψ*, which vanishes with increasing sample size *N*. In general, the estimator overestimates the true parameter. The reason for the overestimate is that the average MOI ranges from 1 to ∞, i.e., the amount by which the parameter can be underestimated is bounded, while it can be arbitrarily overestimated. The non-linear behaviour has the following reason. For small *λ*, occasionally samples with two or more different lineages present are over-represented, and hence the true parameter is greatly overestimated.

**Fig 2.**
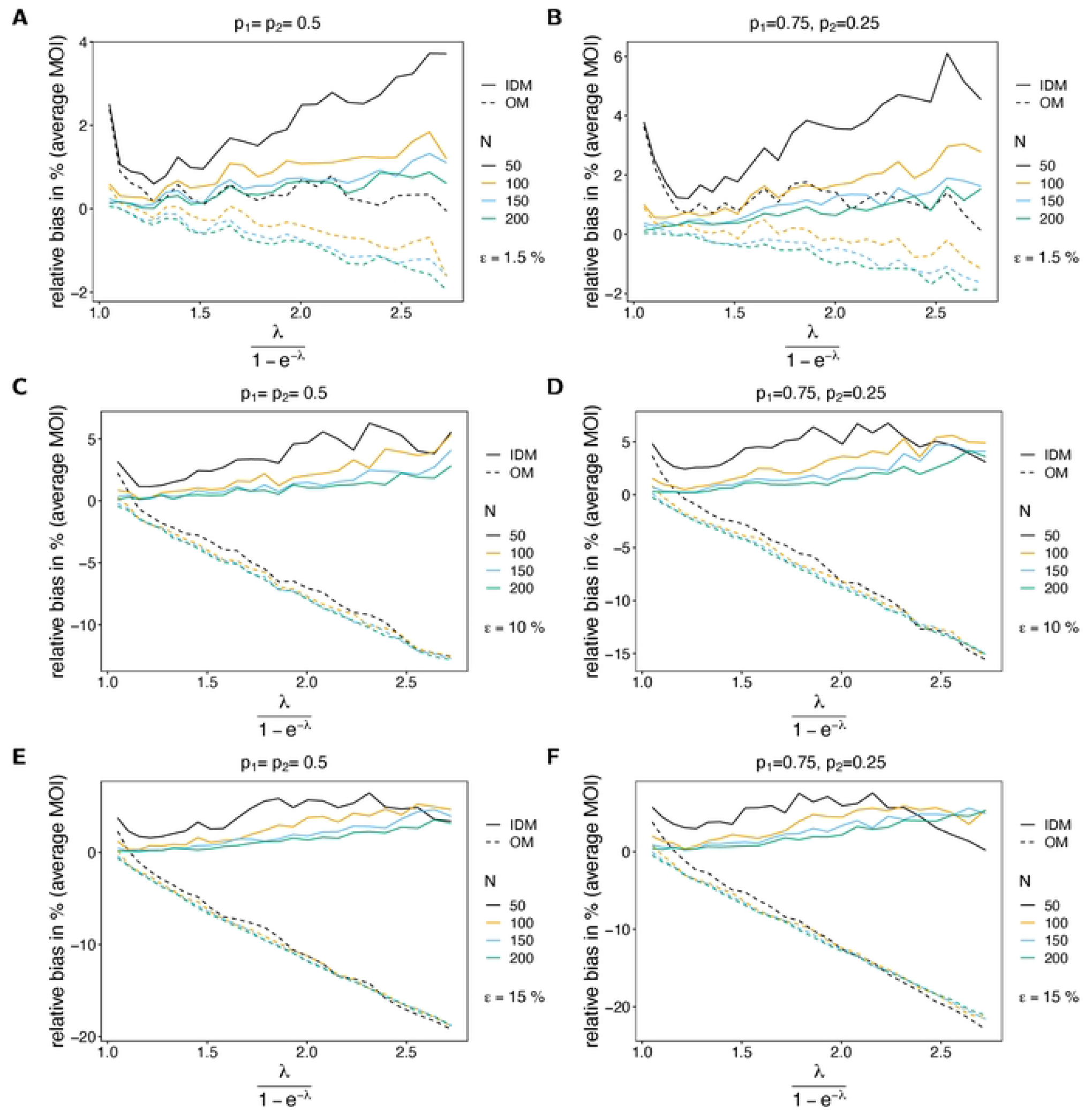
Bias of average MOI. Each panel illustrates the bias of the average 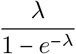 in % of the true average MOI (relative bias) for different sample sizes *N* (colors). Solid lines and dashed lines show the bias of the IDM and OM, respectively. In all panels two lineages (*n* 2) are assumed. Different panels correspond to different probabilities of missing information (*ε*) and frequency distributions (***p***). The left and right panels compare the bias for balanced and unbalanced frequency distributions for the same values of *ε*. In particular, *ε*= 0.015 in (A, B), *ε* =0.1 in (C, D), and *ε* =0.15 in (E, F).

**Fig 3.**
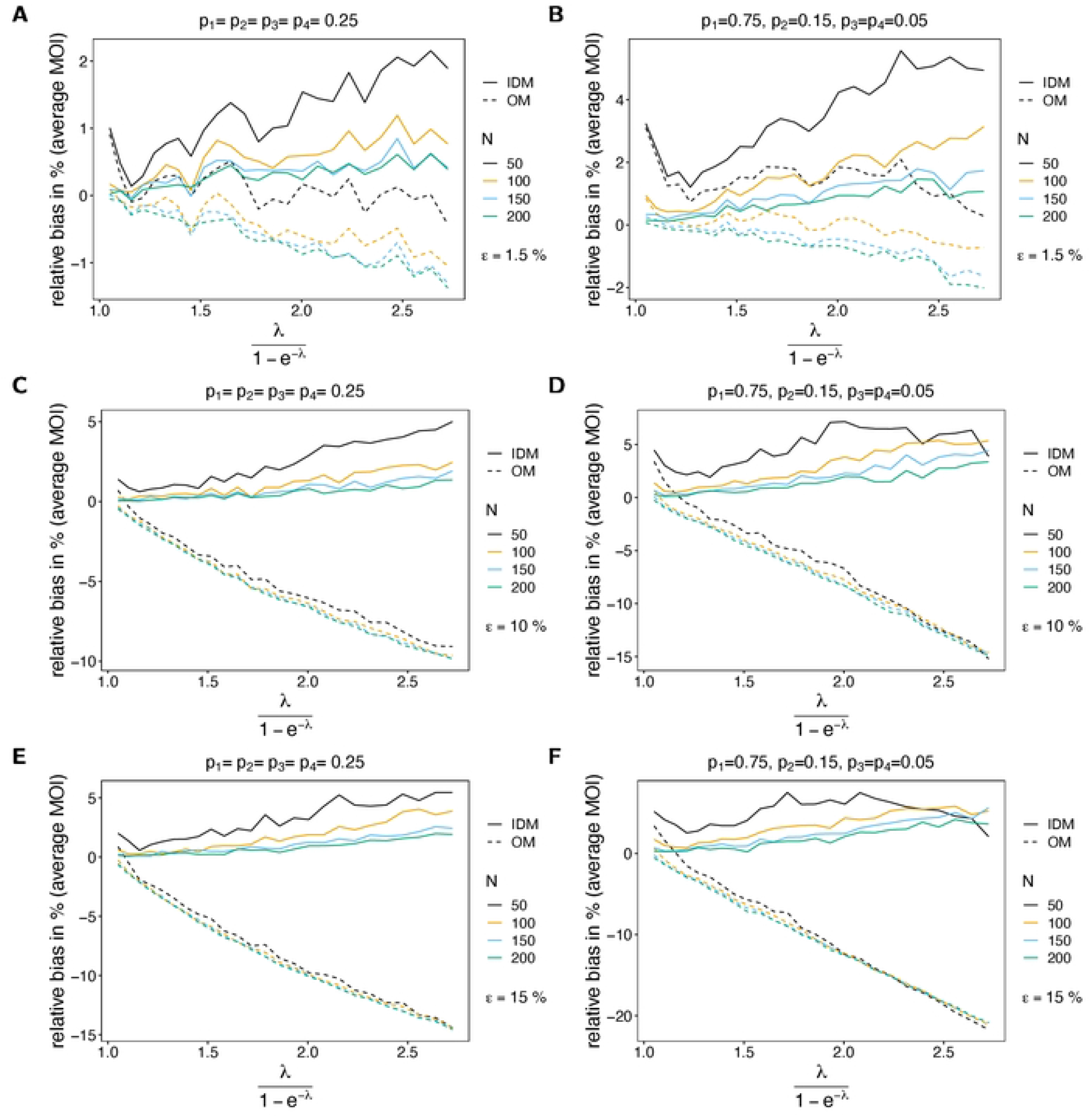
Bias of average MOI. See Fig 2 but for *n* = 4 lineages.

This is typically for MLEs, which are in general sensitive to outliers (cf. Section 5.5 in [17]). (This is particularly true for small sample sizes, where outliers have a stronger effect.) As the true *λ* increases bias starts to decrease and is lowest for values of *λ* corresponding to intermediate transmission. For larger *λ* MOI within samples varies substantially due to the underlying Poisson distribution (which has equal mean and variance *λ*). Consequently, if samples with high MOI are over-represented in a dataset, the MOI parameter is also overestimated. In general, bias is larger for underlying unbalanced frequency distributions (cf. Figs 2, 3A with B, C with D and E with F). The reason is that with unbalanced frequency distributions, it is unlikely to observe infections with different lineages being present unless MOI is high. Hence, the method becomes more sensitive to outliers. In general, the bias is lower with an increase in number of lineages (cf. Fig 2 with Fig 3). The reason is that there is a better resolution to represent super-infections.

If MOI is estimated by the OM, bias behaves quite differently. First, bias substantially increases with an increasing probability of undetected lineages. Particularly the change in bias is strikingly high for large sample size, even if the probability of lineages remaining undetected is as low as *ε*= 1.5%. Second, true parameter is underestimated, except for very small true *λ* (or equivalently *ψ*). Third, bias increases with increasing sample size (*N*). Forth, bias almost linearly decreases as a function of the true parameter *ψ*. The reason is, that many lineages are not observed, which is not reflected at all by the model. This effect is stronger for unbalanced frequency distributions, because (i) infections with different lineages are rare, and (ii) due to incomplete data, it is likely that some of the lineages remain undetected. Hence, observations with no evidence of multiple infections become increasingly likely.

#### Bias of lineage frequency estimates

For the IDM the lineage frequency estimates have very little bias, irrespective of the (i) probability of lineages remaining undetected (*ε*), (ii) sample size, (iii) MOI parameter *λ* (*ψ*), (iv) the number of lineages (*n*) and typically less than 1% (see Fig 4). For symmetric frequency distributions ***p*** the linage frequencies can be considered unbiased. For unbalanced distributions, bias of the dominant lineages seems to increase linearly as a function of the parameter *ψ*. This is not surprising, since in the symmetric case all lineages are equally abundant, whereas in the case of unbalanced frequencies, some lineages are over-represented. Note that bias in Fig 4 fluctuates due to the limited amount of simulation runs. Note further that the bias of minor lineage frequencies in relative terms can be much higher than for predominant lineages, but will be much smaller in absolute terms.

**Fig 4.**
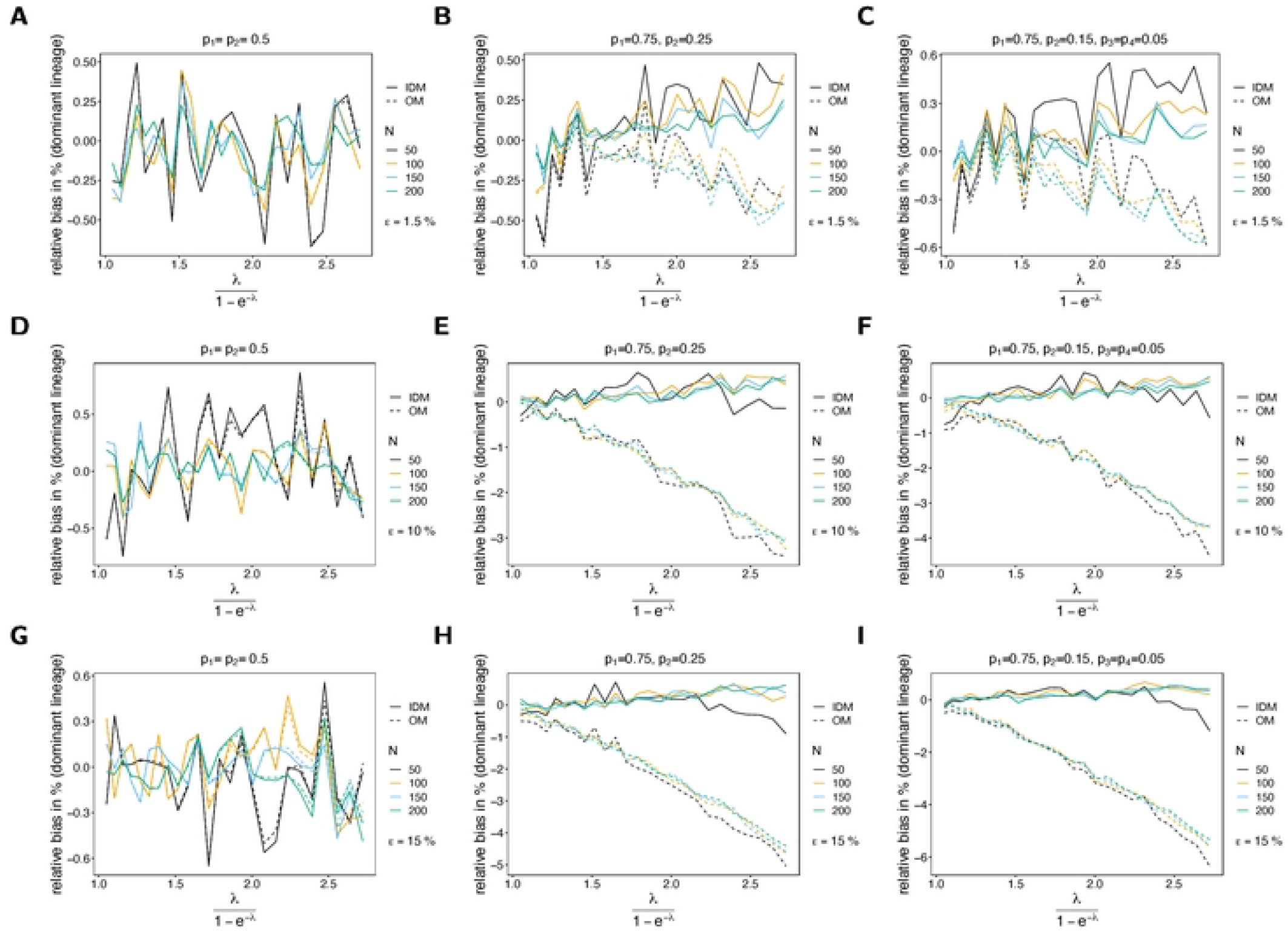
Bias of lineage frequencies. Similar to Fig 2 but for the bias of the dominant lineage in % of the true value as a function of the true average MOI (relative bias). Different panels correspond to different probabilities of missing information (*ε*) and frequency distributions (***p***). While the left panels (A, D, G) assume a balanced frequency distribution, the middle (B, E, H) and the right (C, F, I) panels assume unbalanced frequency distributions.

The bias of lineage frequencies based on the OM changes. Not surprisingly, for symmetric distributions lineage frequencies are unbiased, because all lineages are equivalent. Bias behaves in the same way as for the IDM (see Fig 4A,D,G). On the contrary, for unbalance frequency distributions the dominant lineage tends to be underestimated. Bias is negative and increases linearly with the average MOI *ψ*. For low transmission, the dominant lineage is unbiased, if the probability of lineages remaining undetected is small. However, the bias increases with increasing probability of lineages remaining undetected if transmission is high. Namely, for low transmission (almost only single infections, MOI=1) lineages remain undetected approximately proportional to their frequencies. If transmission is high, the prevalence of rare lineages is high, and they tend to co-occur with predominant lineages. In such infections, both predominant and rare lineages have the same probability to remain undetected, hence leading to an under-representation of the predominant lineages in the data.

#### Bias of the probability of lineages to remain undetected

The probability of lineages to remain undetected is only estimated by the IDM. In general the parameter estimates have little bias, irrespective of the other parameters (Fig 5). The parameter tends to be less biased for larger true parameters *ε*. This is intuitive, as it is estimated to be 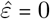 in the absence of empty records, which is very likely if *ε* is small. For unbalanced frequency distributions bias tends to be smaller because empty records are more likely, leading to better estimates of the true parameter.

**Fig 5.**
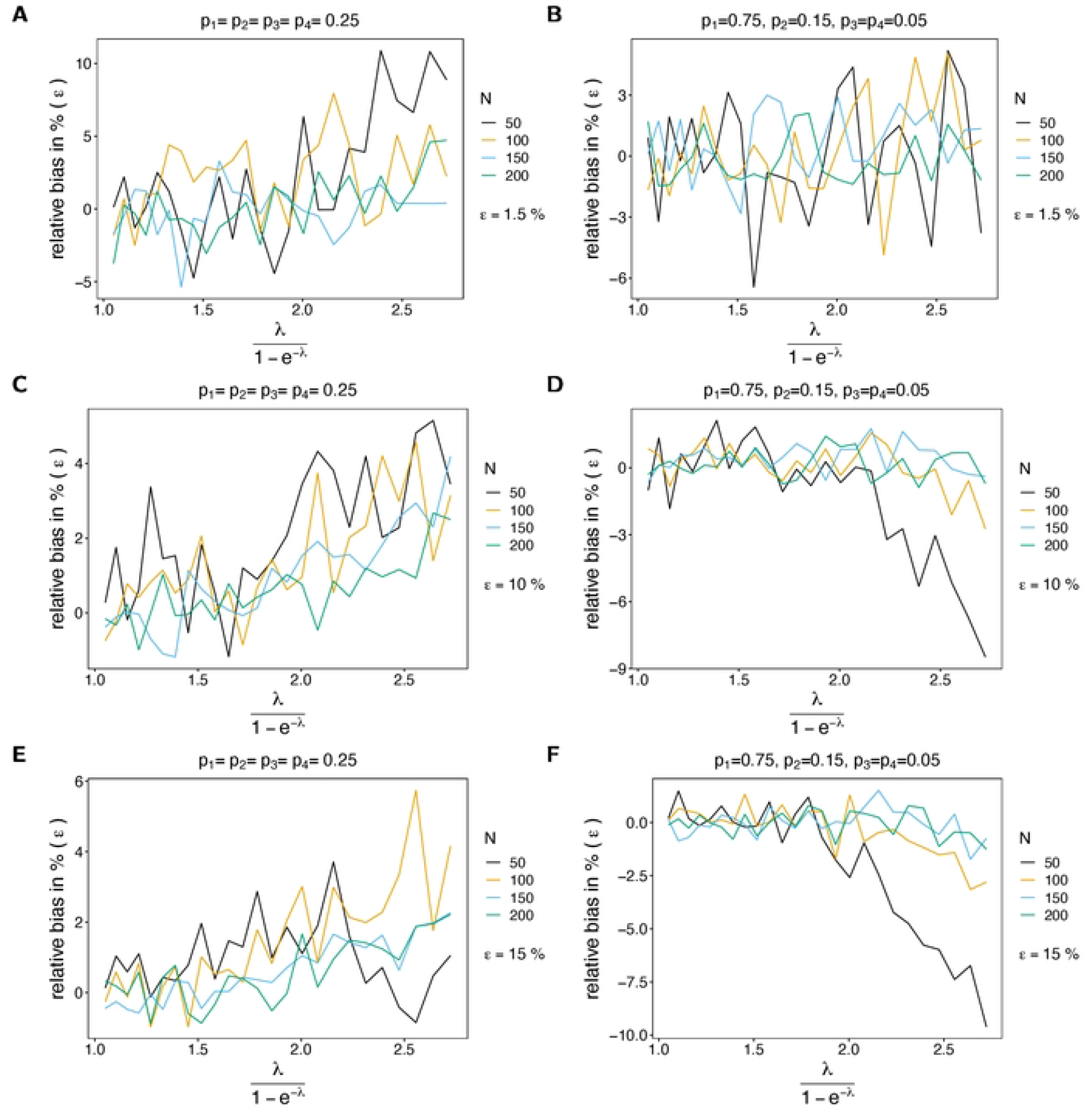
Bias of the probability of lineages to remain undetected. Similar to Fig 2, but for the relative bias of the probability of lineages to remain undetected (*ε*). This parameter is only estimated by the IDM.

#### Variance of the estimator of average MOI

The variance of the estimator based on the IDM in terms of the CV increases as a function of the true parameter. The reason is that for small true *λ* (or *ψ*) datasets tend to be more homogeneous, while the diversity of possible datasets increases with *λ*. This is particularly true for larger numbers of circulating lineages (larger *n*; cf. Fig 6 with Fig 7). The CV of the estimator based on the IDM increases substantially with the likelihood of incomplete observations (larger *ε*). The reason is the following. If no empty records are observed in a dataset, the IDM gives the same estimate as the OM, which tends to be an underestimate (cf. Bias of the average MOI). Particularly, if *ε* is large, the underestimate is substantial. However, once empty records occur in the dataset, the IDM tends to overestimate the true parameter. This creates the large variance of the estimator. Not surprisingly the CV decreases with increasing sample size.

**Fig 6.**
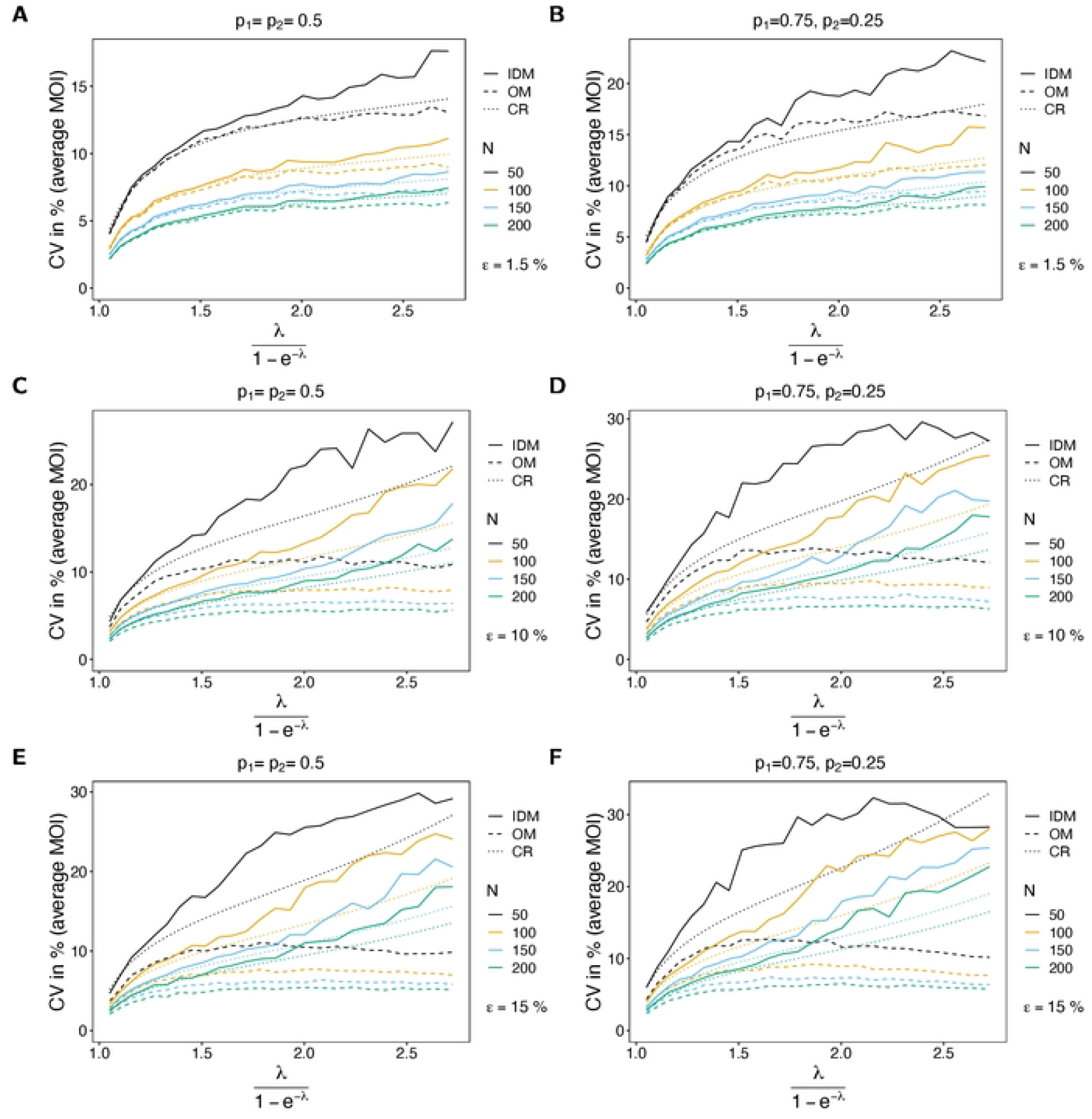
CV of average MOI. Similar to Fig 2, but for the CV of the average MOI in %. The dotted line shows the CV based on the Cramér-Rao lower bound, i.e., the CV is estimated as the square-root of the Cramér-Rao lower, divided by the true parameter.

**Fig 7.**
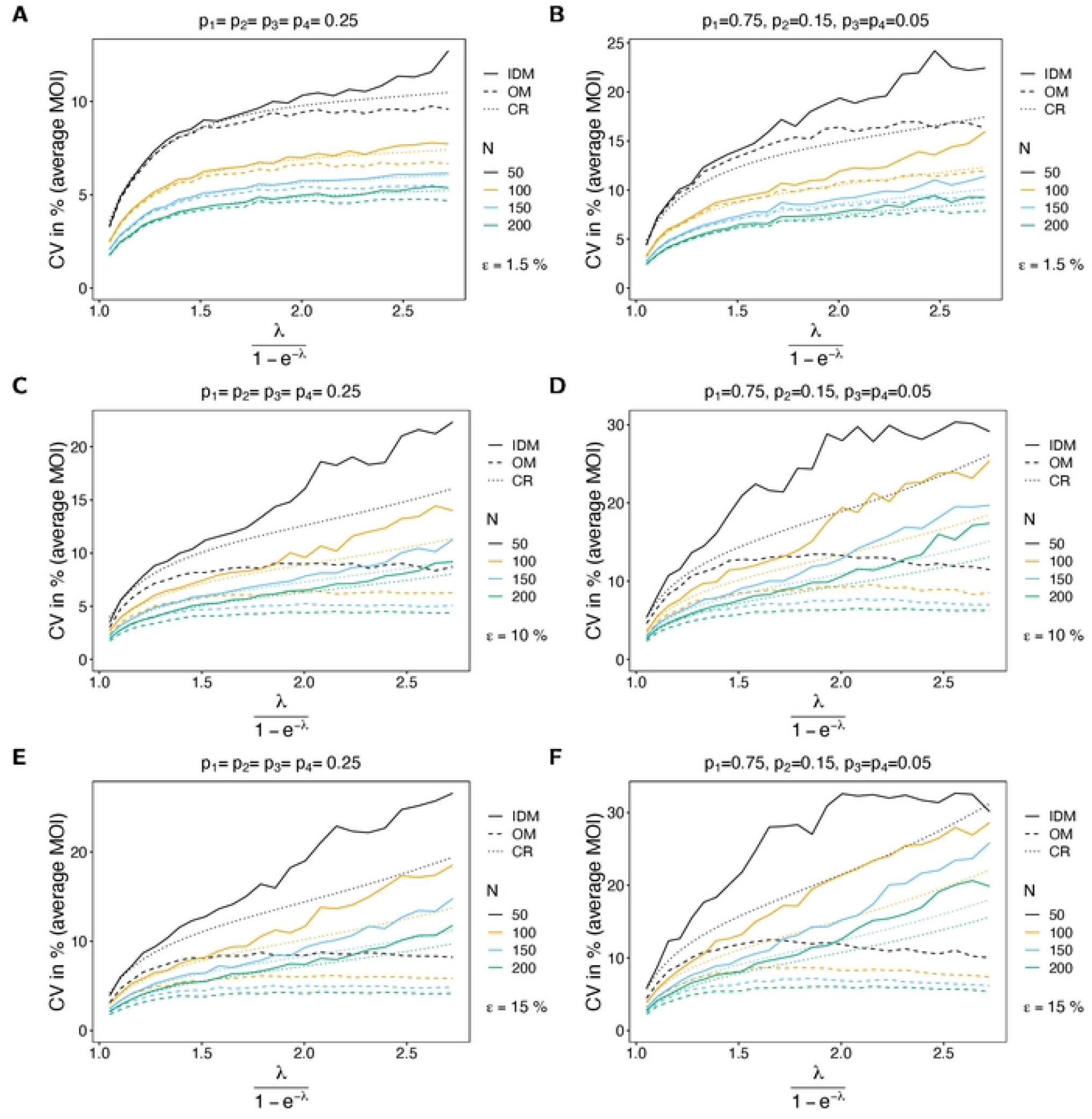
CV of average MOI. See Fig 6 but for *n* = 4 lineages.

For the OM the CV still increases with *ε*, but to a smaller extent, because the parameter is constantly underestimated, which leads to a generally smaller variance. If *ε* increases, empty records effectively lead to a depletion of sample size, which is associated with higher variance.

For small *ψ* the CV almost coincides with the Cramér-Rao lower bound, whereas the CV tends to lie above the lower bound as *ψ* increases (see Fig 6). However, the discrepancy improves for larger sample size *N*. On the contrary, the CV of the estimator based on the OM is lower than the Cramér-Rao lower bound, underlining that this estimator is biased. As expected this becomes more pronounced as *ε* increases.

#### Variance of lineage frequency estimates

Not surprisingly the CV of the lineage frequencies decreases with increasing sample size, because datasets are more informative, leading to more accurate estimates, and hence less variation. Importantly, variance remains is hardly affected by the average MOI *ψ*. However, particularly for balanced frequency distributions, variance tends to be higher for lower transmission, because each lineage has lower prevalence, corresponding to less informative data and hence more variable estimates. This effect is stronger for symmetric frequency distributions (cf. Figs 8A,C,E with B,D,F).

**Fig 8.**
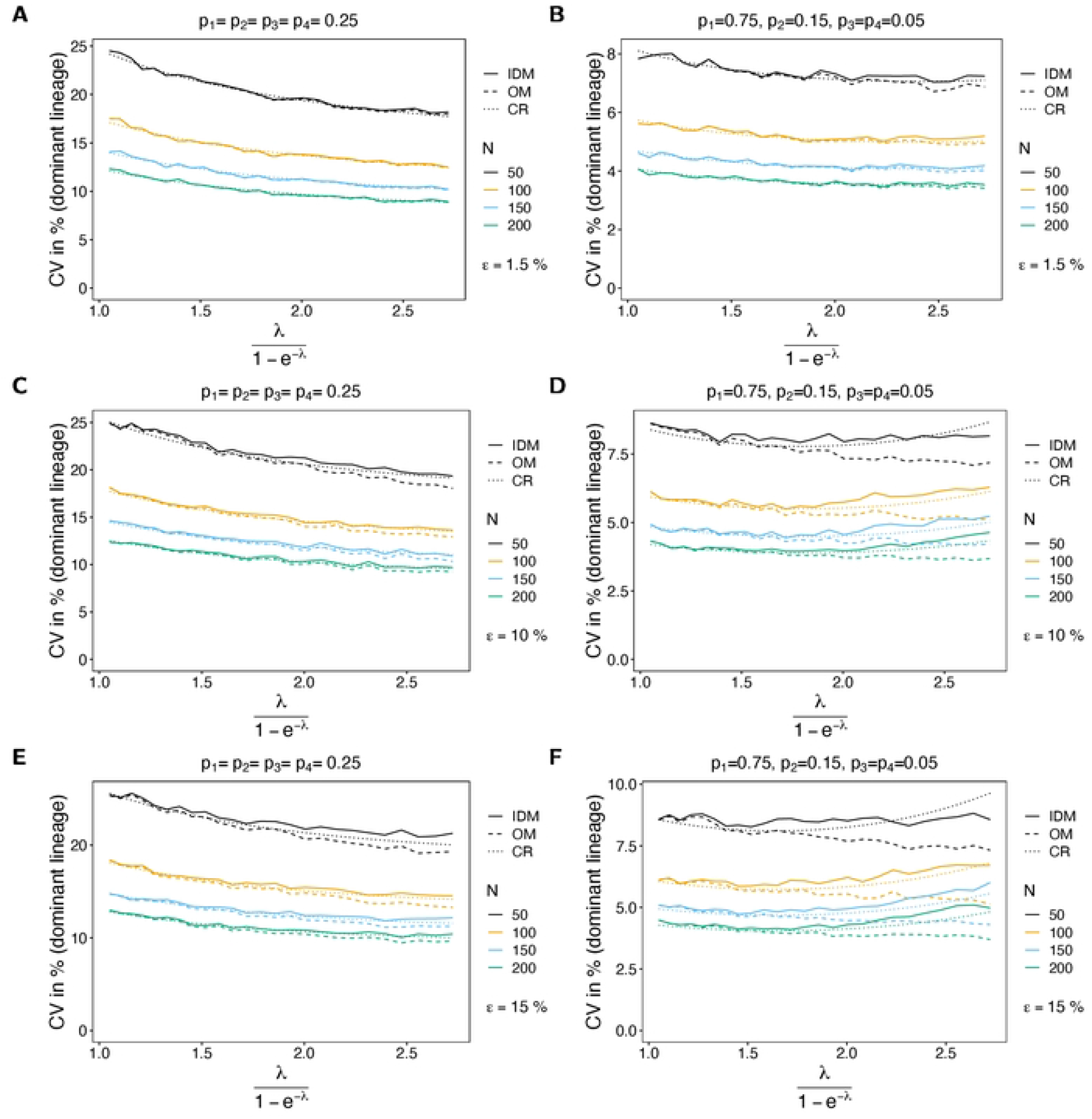
CV of lineage frequencies. Similar to Fig 6 but for the CV of the dominant lineage frequency (*p*_1_) in % as a function of the true average MOI (*ψ*).

The CV is similar for the estimates based on the IDM and OM, but it tends to be slightly lower for the OM. Again there is a god agreement of the CV of the IDM with the Cramér-Rao lower bound.

#### Variance of the probability of lineages to remain undetected

The CV of the probability of lineages to remain undetected is high in general. The reason is that the parameter is estimated as 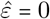 in the absence of empty records, whereas it has a more meaningful estimates if empty records are observed. Consequently, the CV decreases with sample size (empty records are more likely to be observed in a dataset) and increases with MOI (which deceases the chance to observe empty records). For symmetric frequency distributions the CV is higher than for unbalanced distributions (cf. Figs 9A,C,E with B,D,F), because it is less likely to observe empty records. Again there is a reasonable agreement between the CV and the Cramér-Rao lower bound. The fit improves as sample size *N* increases. For small sample size the CV tends to be higher than the Cramér-Rao lower bound, which is not surprising since estimator is sensitive to the number of empty-records, which varies substantially for small sample size.

**Fig 9.** CV of the probability of lineages to remain undetected. Similar to Fig 6, but for the CV of the probability of lineages to remain undetected (*ε*). This parameter is only estimated by the IDM.

#### Mean-squared error of the average MOI estimate

When considering bias and CV of the MOI parameter, it is inconclusive whether the IDM or the OM is preferable. Whereas the estimate of *λ* based on the IDM has lower bias, it has higher variance. Therefore, the mean-squared error (MSE) should be considered (Fig 10). Also no clear preference emerges from the MSE. The MSE of the IDM tends to be lower for large sample size and higher probability of lineages remaining undetected. This is particularly true for balanced frequency distributions (cf. Figs 10A,C,E with B,D,F). If *ε* is small, both models perform similarly well. However, if *ε* is large, the IDM performs only better for larger sample sizes (*N* ≥150).

**Fig 10.** MSE of average MOI. Similar to Fig 2 but for the normalized-root-mean-squared-error of the average MOI in % as a function of the true average MOI (*ψ*).

## Discussion

Because it became feasible to generate molecular data, molecular surveillance of infectious diseases has become increasingly popular [18]. It is particularly useful to reconstruct evolutionary processes of pathogens and routes of transmission, i.e., information that cannot directly be captured from epidemiological data. Especially, for diseases which are endemic and not considered a sudden health emergency, for which asymptomatic infections are common, and/or vector-borne diseases, tracing transmission events directly is almost impossible. Malaria is a typical example of a disease, for which the benefits of molecular surveillance are apparent because only limited information can be retrieved from pure epidemiological surveillance [19]. In fact, some of the most urgent problems in malaria have genetic causes or need the fine-grained information, which can only be mined from genetic/molecular data. This includes monitoring genetic markers associated with drug-resistance, variants associated with failing diagnostics (e.g. HRP2/3 gene deletions in *P. falciparum* causing false-negative rapid diagnostic tests), the distinction between recrudescence and relapses, and transmission intensities [1]. In the context of malaria elimination, monitoring routes of transmission is also highly relevant. Concerning transmission, MOI is an important quantity, which scales with transmission intensities and which is commonly monitored in malaria [2, 19].

Molecular surveillance requires the development of statistical methods adequate to analyze the data being generated. In this context it is also important to address the quality of the molecular data. Namely, molecular assays are not perfect and might lead to missing or erroneous data. A typical problem is the failure of molecular assays, leading to missing data values. This can be neglected as long as missing values can be assumed to be at random. However, missing data values lead to a deflated sample size, which decreases the statistical power of the analysis. On the contrary, if there is a systematic component to missing data, inappropriately accounting for these factors might introduce a bias, which substantially affects the validity and reliability of the results. In the context of MOI - defined here as the number of super-infections (cf. [2]) - there is an intrinsic systematic component to missing information: namely, MOI estimates are based on the presence of genetically distinct variants, observable in clinical specimens (where the presence of identical lineages is reconstructed by a statistical model); if some or all of the distinct variants in samples are frequently not captured by the molecular assay, MOI will be systematically underestimated. If there is reason to assume that data quality is poor due to missing information, it is important to capture this in statistical models.

Here, we introduced a statistical model to estimate the distribution of MOI and the lineage frequency distribution of a single molecular marker. In practice such a marker can be a SNP, STR, or micro haplotype. Such methods are commonly used in malaria molecular surveillance [19]. Estimating the frequency distribution of single markers is often preferential to full haplotype-based methods. Particularly, if one is interested only in the marginal distribution at molecular markers or if the number of possible haplotypes is infeasible.

The proposed model explicitly addresses incomplete information, i.e., with some probability, a lineage remains undetected in a sample. Notably, this is not the only way of modeling incomplete information. An alternative possibility would be to assume that the probability that a lineage remains undetected is proportional to its relative abundance or importance in an infection, reflected, e.g., by the number of times a lineage was super-infecting. Our assumption is based on the following reasons: (i) the abundance of a lineage in a clinical specimen is not necessarily proportional to its abundance within the infection, (ii) the molecular assays, e.g., PCR step, might alter the relative abundance of lineages present in a clinical specimen, and (iii) host-pathogen interactions or the order in which pathogen variants are super-infecting might affect their abundance within the infection. Without more detailed knowledge about host-pathogen interactions and the nature of the molecular assays that produced data, it is not possible to decide for the correct model.

We showed that the proposed model can be written as an exponential family. This guarantees that - except for pathological cases - the maximum-likelihood estimator is well defined, asymptotically unbiased, efficient, and consistent. The pathological cases correspond to the situation that the data lies on the boundary of the admissible data space. We fully classified these cases.

Unfortunately no explicit solution exists for the maximum-likelihood estimate (MLE), so it has to be determined numerically. Here, the EM-algorithm was used to derive a numerical iteration that reliably finds the MLE. The EM-algorithm was derived here and numerically implemented in R. The advantage of this algorithm is its insensitivity to initial conditions, which allows a stable numerical implementation.

We also derived an expression for the prevalence of pathogen variants. From a clinical point of view the prevalence of a pathogen variant (lineage), i.e., the probability that it occurs in an infection, is more relevant than its frequency, i.e., its relative abundance in the pathogen population. Ignoring incomplete information would lead to an underestimation of prevalence. However, appropriate estimates are important when trying to assess, e.g., the prevalence of drug-resistant pathogen variants, as treatment policies will be based on such estimates. Therefore, it was important to provide prevalence estimates accounting for incomplete molecular data.

In the most extreme case all lineages present in a sample fail to be detected (empty records). Usual models to estimate MOI (e.g. [3, 5, 9]) disregard such empty records and assume missingness at random. However, these methods would not account for incomplete information in the remaining observations. This leads to an underestimation of MOI. If empty records do not occur, the proposed estimates coincide with the estimates of the model of [9], which ignores incomplete information. The reason is that the data provides no evidence of incomplete data. Note however, if there is prior information on the probability of lineages to remain undetected, the corresponding model parameter can be set fixed to this prior estimate.

To determine whether the proposed model is an improvement compared to the original model of [3, 5, 9], we conducted a numerical study, to compare both methods performance in terms of bias and variance. It turned out that it is not straightforward which model is superior. Estimates of lineage frequencies tend to be better for the incomplete data model (IDM) if frequency distributions are unbalanced. However, the estimates substantially differ for the MOI parameter. For negligible probabilities of lineages to remain undetected, both models yield similar MOI estimates, in fact they become equivalent in the eliminating case *ε* →0. There is a tendency that the IDM yields less biased MOI estimates than the original model (OM), particularly for larger sample size. However, its coefficient of variation is larger. The larger variation is a result of the nature of the IDM. In the absence of empty records, the MOI estimate is equivalent to that of the OM, and tends to be underestimated. If empty records occur, the MOI parameter tends to be overestimated. On average this creates the larger variance of the estimator. For small probabilities of lineages remaining undetected the mean-squared error (MSE) of the MOI estimate of the IDM is larger than that of the OM. Hence, it is preferable to use the OM. However, if the probability of lineages remaining undetected is large, the estimates of the IDM have a lower MSE. Hence, this model is preferable in such situations. Notably, the fact that the variance of the estimator based on the OM, is smaller than the Cramér-Rao lower bound (which determines the minimum variance of an unbiased estimator), underlines that this estimator is biased.

In conclusion, if there is evidence that the data quality is poor and lineages remain frequently undetected, the proposed model is preferential compared to models that do not appropriately address incomplete information.

## Supporting information

**S1 Mathematical appendix**

**S2 R-script**

**S3 Description of R-script**

## References

1. Organization WH. Global genomic surveillance strategy for pathogens with pandemic and epidemic potential, 2022–2032. World Health Organization; 2022.

2. Schneider KA, Salas CJ. Evolutionary genetics of malaria. Frontiers in Genetics. 2022;13. doi:10.3389/fgene.2022.1030463.

3. Hashemi M, Schneider KA. Bias-corrected maximum-likelihood estimation of multiplicity of infection and lineage frequencies. PLOS ONE. 2022;16(12):1–28. doi:10.1371/journal.pone.0261889.

4. Schneider KA. Large and finite sample properties of a maximum-likelihood estimator for multiplicity of infection. PLOS ONE. 2018;13(4):1–21. doi:10.1371/journal.pone.0194148.

5. Schneider KA, Tsoungui Obama HCJ, Kamanga G, Kayanula L, Adil Mahmoud Yousif N. The many definitions of multiplicity of infection. Frontiers in Epidemiology. 2022;2. doi:10.3389/fepid.2022.961593.

6. Hill WG, Babiker HA. Estimation of Numbers of Malaria Clones in Blood Samples. Proceedings of the Royal Society of London Series B: Biological Sciences. 1995;262(1365):249–257. doi:10.1098/rspb.1995.0203.

7. Davison AC. Statistical Models. Cambridge University Press; 2003. Available from: https://www.cambridge.org/core/product/identifier/9780511815850/type/book.

8. Cohen JE. The biomathematics of malaria. The biomathematics of diseases NO. 1, Norman T. J. Bailey, Charles Griffin and Co., London, 1982. No of pages: xii + 210. Price: £ 16.50, $88.00. Statistics in Medicine. 1984;3(1):93–95. doi:10.1002/sim.4780030111.

9. Schneider KA, Escalante AA. A Likelihood Approach to Estimate the Number of Co-Infections. PLoS ONE. 2014;9(7):e97899.

10. Ihaka R, Gentleman R. R: A Language for Data Analysis and Graphics. Journal of Computational and Graphical Statistics. 1996;5(3):299–314.

11. Bernstein DS. Matrix Mathematics: Theory, Facts, and Formulas with Application to Linear Systems Theory; 2005. Available from: https://www.biblio.com/book/matrix-mathematics-theory-facts-formulas-application/d/1397827927.

12. Pinheiro JC, Bates DM. Mixed-effects models in S and S-PLUS. New York, NY [u.a.]: Springer; 2000. Available from: http://www.worldcat.org/search?qt=worldcat_org_all&q=1441903178.

13. Sundberg R. Statistical Modelling by Exponential Families. Institute of Mathematical Statistics Textbooks. Cambridge University Press; 2019.

14. Davison AC. Statistical Models. Cambridge Series in Statistical and Probabilistic Mathematics. Cambridge University Press; 2003.

15. Barndorff-Nielsen O. Information and exponential families in statistical theory. John Wiley & Sons Ltd; 1978.

16. McLachlan G, Krishnan T. The EM algorithm and extensions. New York: Wiley; 1997.

17. van der Vaart AW. Asymptotic Statistics. Cambridge University Press; 1998. Available from: https://www.cambridge.org/core/product/identifier/9780511802256/type/book.

18. Eybpoosh S, Haghdoost AA, Mostafavi E, Bahrampour A, Azadmanesh K, Zolala F. Molecular epidemiology of infectious diseases. Electron Physician. 2017;9(8):5149–5158. doi:10.19082/5149.

19. Sinha A, Kar S, Deora N, Dash M, Tiwari A, Kori L, et al. India-EMBO Lecture Course: understanding malaria from molecular epidemiology, population genetics, and evolutionary perspectives. Trends in Parasitology. 2023;39(5):307–313. doi:https://doi.org/10.1016/j.pt.2023.02.010.

